# High-resolution glucose tracing and in situ imaging reveals regulated lactate production in human pancreatic β cells

**DOI:** 10.1101/2022.12.21.521364

**Authors:** Federica Cuozzo, Katrina Viloria, Ali H. Shilleh, Daniela Nasteska, Charlotte Frazer-Morris, Zicong Jiao, Adam Boufersaoui, Bryan Marzullo, Hannah R. Smith, Caroline Bonner, Julie Kerr-Conte, Francois Pattou, Rita Nano, Lorenzo Piemonti, Paul R.V. Johnson, Rebecca Spiers, Jennie Roberts, Gareth G. Lavery, Anne Clark, Carlo D.L. Ceresa, Leanne Hodson, Amy P. Davies, Guy. A. Rutter, Masaya Oshima, Raphaël Scharfmann, Matthew J. Merrins, Ildem Akerman, Daniel A. Tennant, Christian Ludwig, David J. Hodson

## Abstract

Using ^13^C_6_ glucose labeling coupled to GC-MS and 2D ^1^H-^13^C HSQC NMR spectroscopy, we have obtained a comparative high-resolution map of glucose fate underpinning β cell function. In both mouse and human islets, the contribution of glucose to the TCA cycle is similar. Pyruvate-fueling of the TCA cycle is primarily mediated by the activity of pyruvate dehydrogenase, with lower flux through pyruvate carboxylase. While conversion of pyruvate to lactate by lactate dehydrogenase (LDH) can be detected in islets of both species, lactate accumulation is six-fold higher in human islets. Human islets express LDH, with low-moderate *LDHA* expression and β cell-specific LDHB expression. LDHB inhibition increases glucose-dependent lactate generation in mouse and human β cells, and decreases Ca^2+^-spiking frequency without affecting ATP/ADP levels. Thus, we show that LDHB limits glucose-stimulated lactate generation in β cells. Further studies are warranted to understand how lactate impacts β cell metabolism and/or function.

**HIGHLIGHTS:**

- Human and rodent islets generate lactate following glucose stimulation.
- β cells specifically express LDHB, which acts to limit lactate generation.
- LDHB inhibition influences Ca^2+^ spiking frequency without affecting ATP/ADP ratio.

**eTOC:** Cuozzo et al show that glucose-stimulated rodent and human islets generate lactate. Transcriptomic and imaging analyses reveal that LDHB is specifically expressed in β cells and unexpectedly restrains lactate production. LDHB expression and thus regulated lactate generation might reflect a key mechanism underlying β cell metabolism, function and survival.

## INTRODUCTION

β cells are highly-adapted as glucose sensors and need to balance glucose-dependent insulin release with housekeeping metabolic functions. The traditional view of β cell metabolism focuses on a tight relationship between blood glucose concentration, oxidative phosphorylation and stimulus-secretion coupling. Following a rise in glycemia, glucose enters the β cell through facilitated transport via low affinity glucose transporters (GLUT1 and GLUT2 in humans and rodents, respectively) ^1,2^. Glucose is then phosphorylated by a low affinity hexokinase, glucokinase (GK), leading to closure of the ATP-sensitive potassium (K_ATP_) channels (reviewed in ^3,4^). The increase in membrane voltage then drives Ca^2+^ flux through voltage-dependent Ca^2+^ channels ^3^, which together with amplifying signals (amino acids, isocitrate, cAMP etc) ^4–6^, drives first and second phase insulin granule exocytosis. Direct conversion of pyruvate to lactate is thought to be suppressed in the β cell due to low levels of lactate dehydrogenase A (*LDHA*) ^7–10^, ensuring that the majority of pyruvate enters the TCA cycle.

Recent studies have challenged the canonical view of β cell metabolism by showing that ATP/ADP generation might be compartmentalized. K_ATP_ channels are locally regulated by a membrane-associated glycolytic metabolon that is assisted by the mitochondrial phosphoenolpyruvate (PEP) cycle ^11–13^. The glucose-dependent rise in the ATP/ADP ratio raises mitochondrial voltage to stall oxidative phosphorylation and the TCA cycle, activating anaplerotic flux through pyruvate carboxylase (PC) and the PEP cycle, which supports ATP production by pyruvate kinase (PK) until K_ATP_ channels are closed. Following membrane depolarization, the rise in ADP supports a highly oxidative state that depends on high TCA cycle flux and pyruvate consumption by pyruvate dehydrogenase (PDH), which supports oxidative phosphorylation and sustained secretion ^11,12,14^. These two complementary states might dictate the mitochondrial fate of pyruvate.

Alongside compartmentalized ATP/ADP generation, anaplerotic metabolism serves as an important source of coupling or amplifying factors for glucose-stimulated insulin secretion ^15–17^. For example, PC directs pyruvate entry into the TCA cycle, which feeds isocitrate into isocitrate dehydrogenase 1 (IDH1) to support 2-ketoglutarate (2-KG)/NADPH generation and SENP1 activation ^6,18^. This reaction is further supported by metabolism of glutamine by the reductive TCA cycle ^19^.

Despite the clear importance of metabolism for β cell insulin release and phenotype, we are still lacking a high-resolution, integrated view of β cell glucose fate. In particular, most data using glucose tracing and GC-MS/NMR spectroscopy has been derived from insulinoma cell lines, which provide the requisite cell mass for metabolite detection/annotation. However, insulinoma cell lines have to balance the need for insulin secretion with proliferation, an energy-consuming process ^15,17,20–23^, and fail to display normal cell heterogeneity known to influence metabolism ^24–26^. In addition, species-differences in islet cell composition and β cell function have been described ^27–29^, yet their influence on energetics is still unclear. Lastly, bulk metabolomics has been informative for understanding β cell metabolism ^30,31^ and, while sensitive, lacks the resolution required to pinpoint glucose fate during glycolysis and the TCA cycle. Recent studies have shown the power of metabolic tracing to compare stem-derived β cell and human islet metabolism, highlighting key differences in glucose metabolism, metabolite trafficking and sensitivity to various fuels ^32,33^. However, these studies did not (understandably) combine GC-MS/LC-MS with NMR for TCA metabolite annotation, nor cross-compare human and mouse islets. Thus, our understanding of β cell glucose metabolism in mouse and human islets remains incomplete.

In the present study, we combine GC-MS-based ^13^C_6_ glucose tracing with the resolution of 2D ^1^H-^13^C HSQC NMR multiplet analysis to map glucose fate in islets with high sensitivity. By applying this dual approach to human and mouse samples, we are able to provide a detailed cross-species depiction of glucose metabolism. Detailed examination of ^13^C labelling patterns confirms that PDH is the major contributor to the TCA cycle. We further show that pyruvate is directly converted to lactate in both human and mouse islets. However, lactate accumulation is much higher in human islets. Analysis of multiple transcriptomic datasets shows that *LDHB* expression is restricted to human β cells, confirmed using immunohistochemistry for LDHB protein. In situ metabolic imaging using specific inhibitors shows that LDHB limits lactate production, but does not affect glucose-stimulated ATP/ADP and Ca^2+^ rises. We thus provide an integrated view of mouse and human islet metabolism, show that β cell-specific LDHB expression limits lactate generation, and suggest that the role of lactate in β cell metabolism, function and survival is worth revisiting.

## RESULTS

### Glucose contribution to TCA cycle in human and mouse islets

To investigate glucose handling, mouse and human islets were incubated overnight with ^13^C_6_ glucose prior to metabolite extraction and GC-MS and 2D ^1^H,^13^C HSQC-NMR spectroscopy (Figure 1A). To allow sufficient glucose flux for detection of ^13^C incorporation into TCA metabolites, without inducing glucotoxicity, 10 mM ^13^C_6_ glucose was used. The incorporation of ^13^C from ^13^C_6_ glucose into the TCA cycle metabolites was then established via mass isotopologues distribution (MID) analysis (Figure 1B).

**Figure 1:**
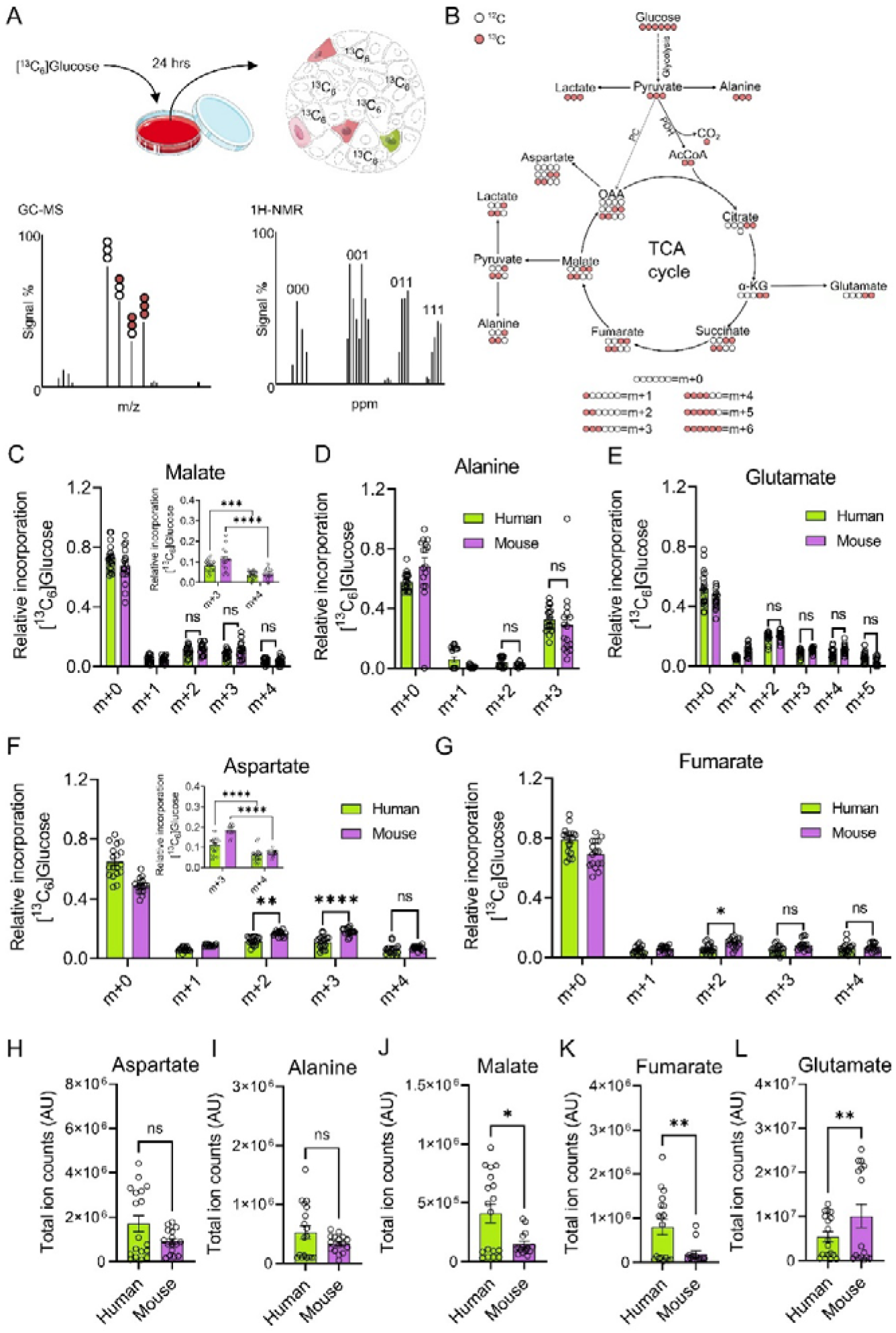
MID analysis of glucose fate in human and mouse islets. **A)** Schematic showing GC-MS and ^1^H-NMR-based ^13^C_6_ glucose-tracing protocol in primary islets. **B)** Schematic showing mass isotopomer distribution (MID) analysis of ^13^C_6_ glucose-tracing data. **C-E)** MID analysis showing similar incorporation of ^13^C from ^13^C_6_ glucose into malate (C), alanine (D) and glutamate (E) in human and mouse islets. **F, G**) MID analysis showing increased incorporation of ^13^C from ^13^C_6_ glucose into m+2 and m+3 aspartate (F), and m+2 fumarate (G) in mouse compared to human islets. **H, I)** Total amount of extracted aspartate (H) and alanine (I) is similar in human and mouse islets. **(J-L)** Total amount of extracted malate (J) and fumarate (K) is decreased in mouse relative to human islets, whereas glutamate (L) is increased. For all data, n = 18 independent replicates from 9 human donors; n = 10 islet preparations from 15 animals. C-G were analyzed using two-way ANOVA and Sidak’s post-hoc test. H-L were analyzed using Welch’s test. Bar graphs show individual datapoints and mean ± SEM. AU = arbitrary unit.

Suggesting a similar progression of glycolysis and the TCA cycle, glucose incorporation into glycerol-3-phosphate (G-3-P) (Figure S1A) and the major metabolites malate, alanine, and glutamate was not different between mouse and human islets (Figure 1C-E). However, a slight but significant increase in m+2/m+3 aspartate and fumarate was detected in mouse islets (Figure 1F and G), reflecting an increased contribution of glucose-derived pyruvate into the TCA cycle via acetyl-CoA. Total aspartate and alanine levels did not differ between the species (Figure 1H and I), whereas malate and fumarate levels were lower in mouse (Figure 1J and K). Glutamate levels were ∼3-fold higher in mouse versus human islets, despite similar MIDs, implying that there is a larger contribution of non-labelled glutamate to the total glutamate pool in this species e.g. through glutamine transport (Figure 1L).

### Lactate generation is higher in human compared to mouse islets

To obtain a higher definition view of pyruvate management, its contribution to the production of alanine and lactate was assessed. In both species, glucose incorporation could be detected in m+2 and m+3 lactate, derived from the TCA cycle and direct pyruvate conversion, respectively (Figure 2A-C). While the MID for alanine was similar in islets from both species (Figure 1D), the accumulation of m+2 and m+3 lactate was significantly (∼ 6-fold) higher in humans (Figure 2A). The total amount of lactate was higher in human than in mouse islets (Figure 2B-D). Supporting the notion of increased lactate production through glycolytic input in human islets, the m+3 G-3-P /m+3 lactate labeling ratio was higher in mouse versus human islets (Figure S1B). While accumulation of m+2 lactate was expected, we were surprised to detect significant m+3 lactate accumulation in human islets, since *Ldha* has been shown to be “disallowed” in rodent β cells, hence preventing alternative fates for pyruvate ^7–10^.

**Figure 2:**
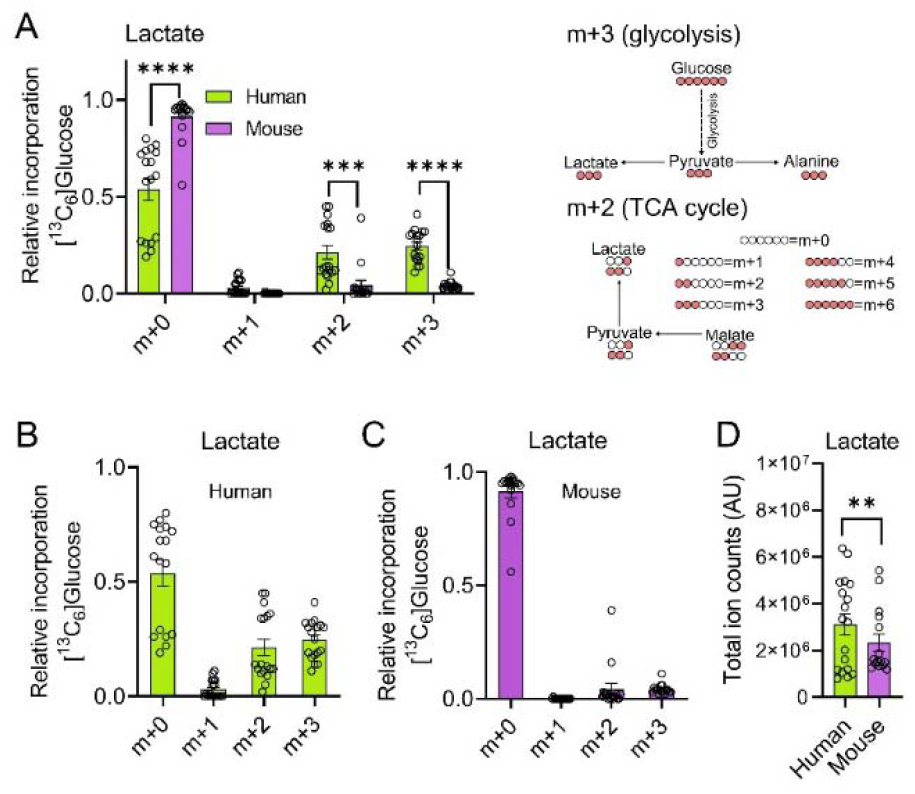
Human islets produce lactate. **A-C)** MID analysis shows detectable glucose incorporation into m+2 (TCA cycle) and m+3 (pyruvate conversion) lactate, with more accumulation in human (A, B) versus mouse (A, C) islets (n = 18 independent replicates from 9 human donors; n = 10 islet preparations from 15 animals) (two-way ANOVA and Sidak’s post-hoc test). **D)** Total lactate generation is higher in human compared to mouse islets (n = 18 independent replicates from 9 human donors; n = 10 islet preparations from 15 animals) (Welch’s test). Bar graphs show individual datapoints and mean ± SEM.

Suggesting that m+3 lactate is derived mainly from pyruvate, rather than multiple rounds of the TCA cycle, were the following findings: 1) m+4 malate and m+4 aspartate were significantly lower compared to m+2 and m+3 isotopomers (Figure 1C and F); and 2) m+4 malate and m + 4 aspartate were similar in human and mouse, despite significantly higher m+3 lactate in human (Figure 1C and F). In addition, previous tracing studies have shown m+3 lactate accumulation in iPSC-derived islets and human islets after 1 hour of tracing ^33^. Lastly, lack of lactate detection in previous studies with mouse islets and rodent beta cell lines is unlikely due to the shorter timecourses used, since we were also unable to observe meaningful m+3 lactate generation after 24 hours incubation with ^13^C_6_-glucose.

### High-resolution annotation of ^13^C_6_ glucose tracing data

To identify isotopomer patterns with high-resolution, the MID analysis of ^13^C_6_ glucose-traced human and mouse islets was annotated with 2D ^1^H-^13^C HSQC NMR multiplet analysis. From uniformly labeled glucose, ^13^C atoms are incorporated into the metabolites of the TCA cycle through the activity of PDH and PC (Figure 3A, B). This leads to the formation of labeling patterns within the chemical structure of each metabolite that are specific to the pathway from which they are produced (Figure 3A, B). Therefore, the positions of ^13^C atoms within each metabolite can be utilized to elucidate the relative activities of PDH and PC. To define the different isotopomer patterns a numerical notation was used, where the numbers 0 and 1 indicate ^12^C and ^13^C atoms, respectively. Confirming the accuracy of the approach and the robustness of our findings, the accumulation of lactate_111_ (i.e. fully-labeled lactate) was significantly higher in human compared to mouse islets, in line with the MID glucose-tracing data (Figure 2A-C) (Figure 3C-E).

**Figure 3:**
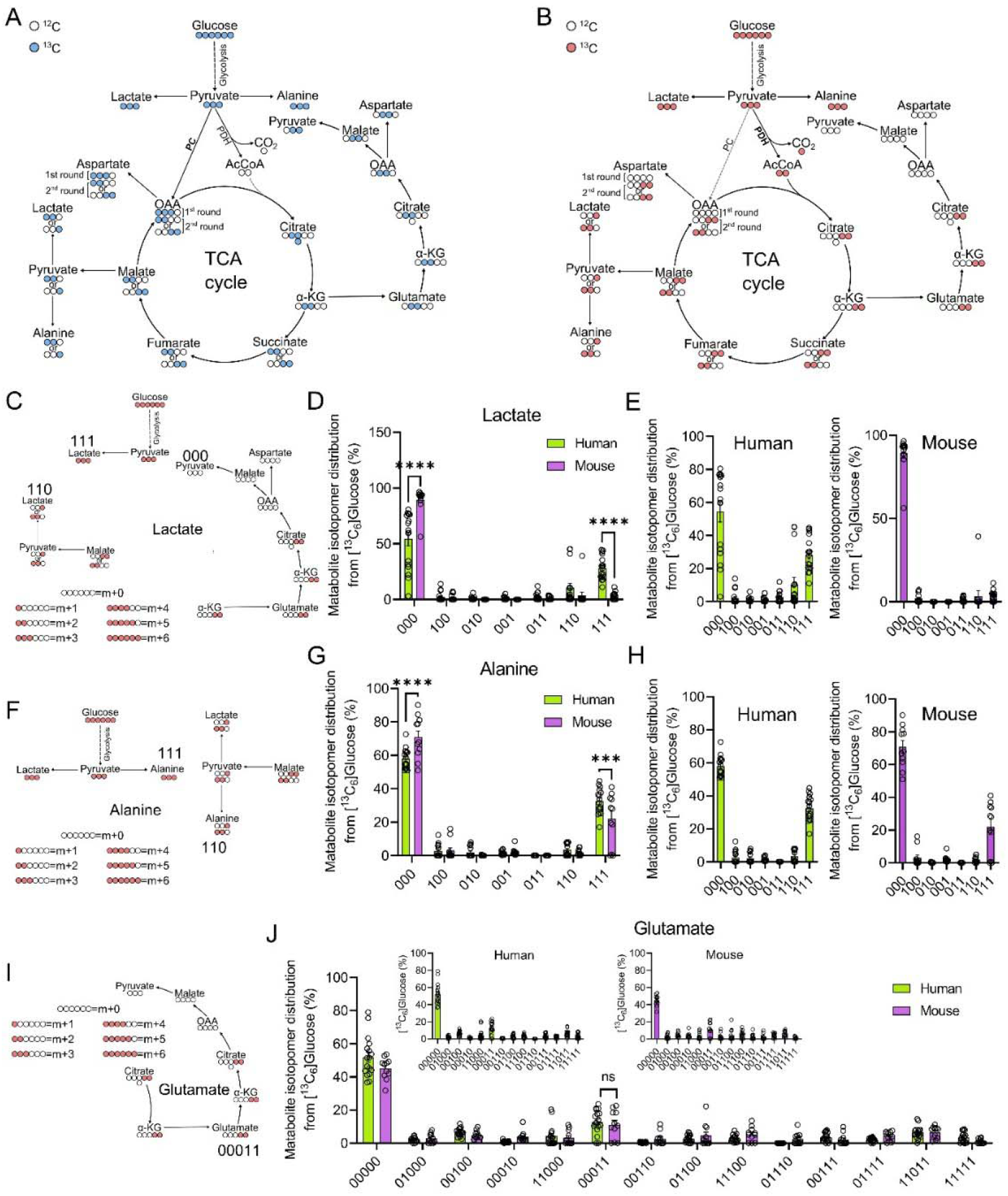
Incorporation of ^13^C from ^13^C_6_ glucose into TCA cycle metabolites through PDH and PC. **A)** White and blue circles, respectively, show the incorporation of ^12^C and ^13^C into TCA cycle metabolites arising from metabolism of pyruvate by PC. **B)** White and red circles, respectively, represent ^12^C and ^13^C atoms as incorporated from ^13^C_6_ glucose into the TCA cycle through the conversion of pyruvate to acetyl-CoA by PDH. **C-E)** Lactate_000_, lactate_111_ and lactate_110_ are the most abundant isotopomers (C) in both humans and mice (D), although the incorporation of ^13^C from ^13^C_6_ glucose into lactate_111_ is significantly higher in human than mouse islets (D, E). **F-H)** ^13^C incorporation into alanine isotopomers (F) is similar in human and mouse islets (G), with alanine_111_ being the most represented labeled isotopomer (G, H). **I, J**) The distribution of labeling patterns for glutamate (I) are similar in human and mouse islets (J), with glutamate_00011_ being the most abundant labeled isotopomer in both species (J). For all data, n = 16-17 islet preparations, 9 human donors and n = 12-15 islet preparations, 7-8 animals. Data were analyzed using two-way ANOVA and Sidak’s post-hoc test. Bar graphs show individual datapoints and mean ± SEM. AU = arbitrary unit.

### TCA cycle fuelling is more dependent on flux through PDH than PC in human and mouse islets

In both human and mouse islets, lactate_110_ made a greater contribution to the m+2 isotopologue pool than the other possible isotopomers (Figure 3C-E). This finding suggests that TCA-derived lactate is produced primarily from the oxidative TCA cycle rather than the reductive metabolism of PC-derived glutamate, from which pyruvate_011_ and then lactate_011_ would arise (Figure 3C-E). We also noticed that the majority of alanine was either 000 or 111, with only a very minor contribution to the other isotopomers (Figure 3F-H). As such, the labeled portion of alanine is exclusively produced from pyruvate upstream of the TCA cycle, meaning that the accumulation of pyruvate_110_ from malate_1100_ (i.e. TCA-derived) is almost entirely employed to regenerate cytoplasmic lactate_110_ (Figure 3F-H). Alanine_111_ accumulation was slightly (∼20%) higher in human than mouse islets, reflecting a greater contribution of transamination toward amino acid production (Figure 3F-H). Supporting the lactate isotopomer data, the contribution of ^13^C_6_ glucose to the labeling patterns of glutamate was found to be similar in humans and mice (Figure 3I, J). In both species, the most abundant labeled isotopomer was glutamate_00011_ (Figure 3I, J), which is derived from TCA cycle flux through the activity of PDH (Figure 3A, J). Together, these findings provide further evidence that pyruvate management in the pancreatic β cell occurs primarily through PDH at the stimulatory glucose concentration used here ^15,34^.

### LDH is expressed at higher levels in human compared to mouse **β** cells

Given that lactate generation was detected in human islets, and to a lesser extent mouse islets, we set out to investigate whether LDH protein is expressed in β cells. To this end, we performed immunohistochemistry using an antibody with cross-reactivity against LDHA, LDHB and LDHC (i.e. total LDH). Since LDHC is undetectable in the pancreas ^35^ and is largely confined to the testis ^36^, we assume that total LDH staining in pancreas represents LDHA and LDHB isoforms. Punctate cytoplasmic LDH staining was detected in human endocrine and exocrine pancreas, including in β cells (Figure4A). LDH expression was much lower in mouse compared to human islets, with slightly higher levels in the exocrine versus endocrine compartments (Figure 4A and C). However, following 8 weeks high-fat diet (HFD) feeding to induce beta cell de-differentiation, LDH protein expression increased ∼ 2-fold versus age-matched standard diet controls (Figure 4B and C). Of note, LDH expression also increased in the exocrine compartment, with HFD mouse pancreata resembling more their human counterpart (Figure 4A-D). Occasional intra-islet cells were found to express very high LDH levels in both human and mouse, likely representing endothelial cells known to be enriched for LDHA ^37^ (correlation for enrichment = 0.502; Human Protein Atlas, proteinatlas.org) (Figure 4A and B). β cell de-differentiation was confirmed in the same samples using immunostaining for PDX1, which was significantly lower under HFD versus standard chow conditions (Figure 4E and F).

**Figure 4:**
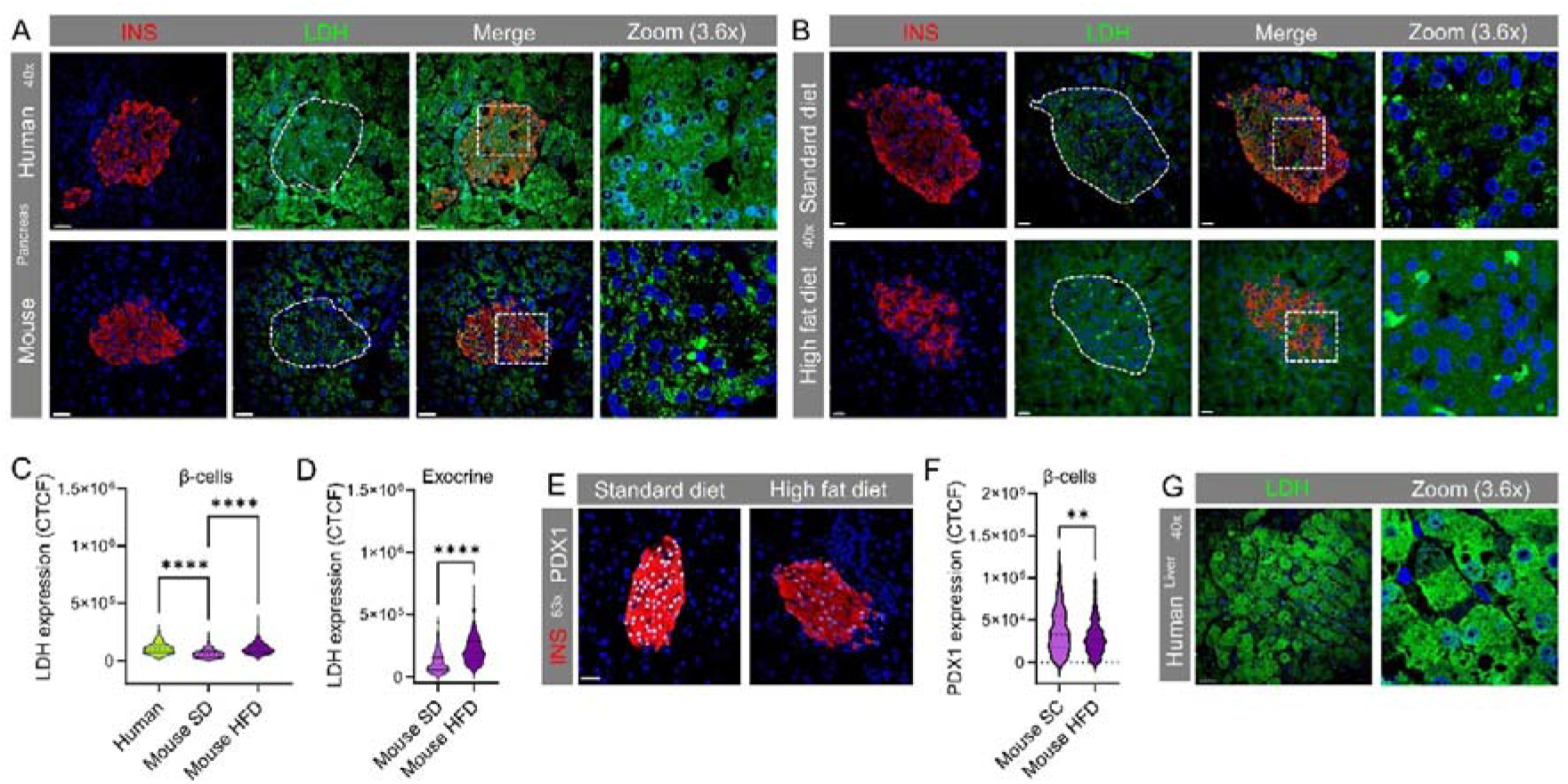
Human islets express LDHA, LDHB and LDHC protein. **A)** LDH expression, assessed using an antibody with cross-reactivity against LDHA, LDHB and LDHC, is higher in human β cells compared to mouse β cells. **B)** LDH expression increases in mouse exocrine tissue and β cells following 8-12 weeks high fat diet (HFD) feeding versus age-matched standard diet (SD) controls. **C)** Corrected total cell fluorescence (CTCF) quantification of LDH protein expression in human β cells (n = 240 cells, 3 donors), SD mouse β-cells (n = 220 cells, 3 animals), and HFD mouse β cells (n = 250 cells, 3 animals) (Kruskal-Wallis test, Dunn’s post-hoc test). **D)** As for G), but showing LDH immunoreactivity in the exocrine pancreas of mice fed either SD or HFD (n = 230-250 cells, 3 animals) (Mann-Whitney test). **E)** PDX1 expression is decreased in β cells of mice fed HFD for 8 weeks versus SD controls (n = 240 cells, n = 3 animals). **F)** CTCF) quantification of PDX1 protein expression in SD and HFD mouse β-cells (n = 240 cells, 3 animals) (Mann-Whitney test). **G)** Immunostaining showing strong LDH expression in human liver from donors with non-alcoholic steatohepatitis (n = 240 cells, n = 3 donors) (Mann-Whitney test). Representative images are shown. Scale bar = 30 µm. Bar graphs show individual datapoints and mean ± SEM. Violin plot shows median and interquartile range. AU = arbitrary unit.

Strong LDH staining could be detected in human liver sections derived from patients with non-alcoholic steatohepatitis (NASH), in which hepatocytes express high LDH levels (correlation for enrichment = 0.502; Human Protein Atlas, proteinatlas.org) (Figure 4G) ^38,39^. As expected from studies of enzyme activity, LDH expression was higher in liver than in β cells (Figure S2A-C) (LDH CTCF mean ± SD: 1.16 x 10^6^ ± 2.06 x 10^5^ versus 2.85 x 10^5^ ± 4.90 x 10^4^, liver versus β cell; P<0.0001, Mann-Whitney test) (n = 240 cells, n = 3 donors) ^8^. All results were confirmed with multiple immunostaining runs, using both 40x and 63x objectives (Figure S2A-D).

### *LDHB* is expressed in human **β** cells within the endocrine compartment

Next, we interrogated published RNA-sequencing studies to understand which LDH isoenzymes might underlie lactate generation in human islets. Using data from FACS-purified fractions, *LDHB* was found to be specifically- and highly-expressed in β cells (shown also in ^40^) within the islet, whereas the opposite convention was observed for *LDHA* (i.e. specifically and highly expressed in α cells) (Figure 5A). Analysis of single cell RNA-seq data confirmed these findings across multiple independent datasets (Figure 5B and C), as visualized according to endocrine and non-endocrine cell clusters (Figure 5D and E). Confirming accuracy of the clustering-based analysis (and inability of β cells to export lactate), the lactate transporter MCT1, encoded by *SLC16A1*, could not be detected in any islet endocrine cell type (Figure 5F), as previously reported ^41,42^.

**Figure 5:**
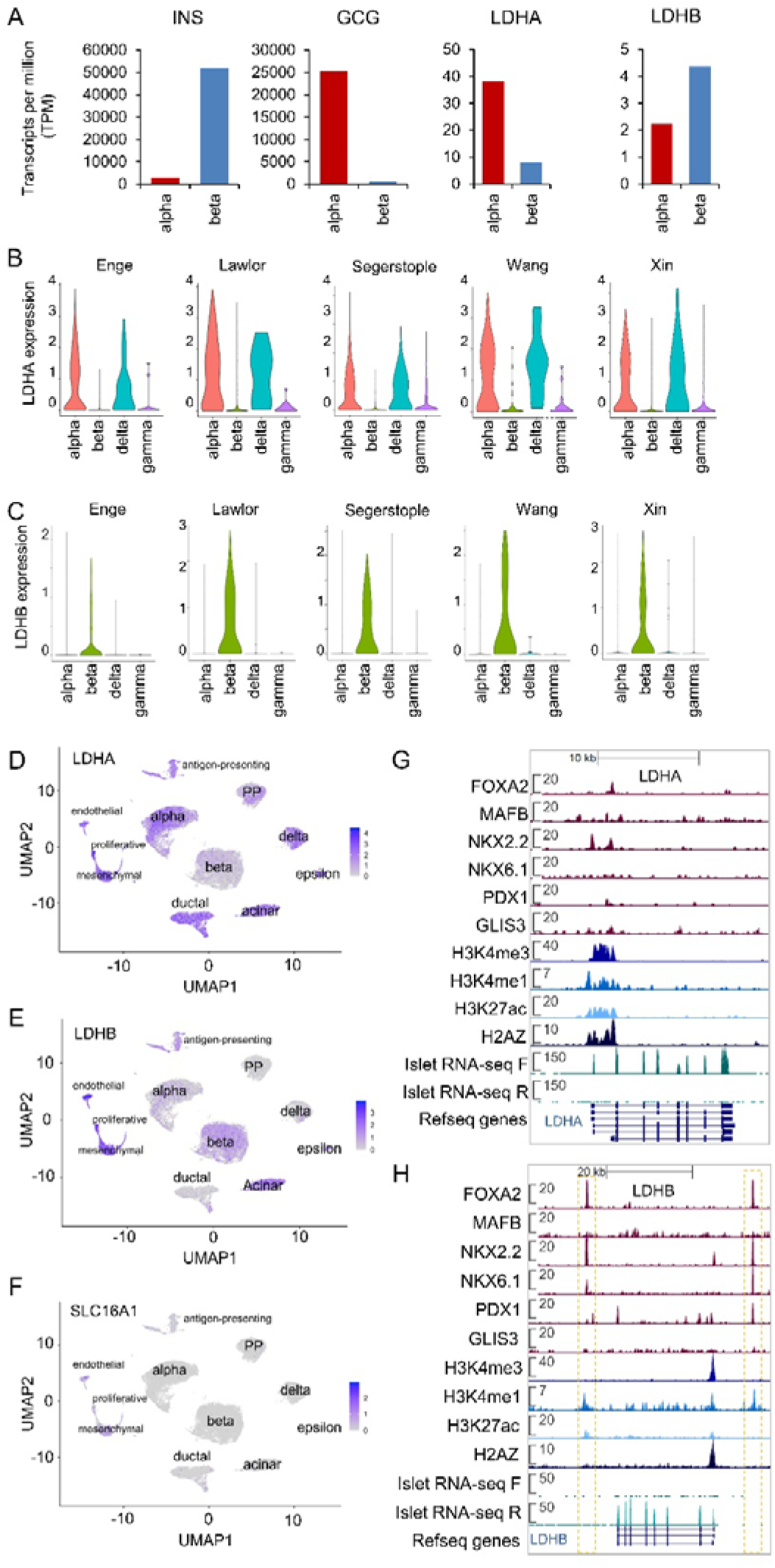
Human β-cells specifically express *LDHB*. **A)** Normalized mRNA levels (transcripts per million, TPM) for *INS*, *GCG*, *LDHA* and *LDHB* genes in fluorescent-activated cell-sorted (FACS) α and β cell samples (re-analysis of data from ^74^). **B, C)** Normalized LDHA (B) and LDHB (C) expression in α, β, δ and γ cells from five independent human islet single cell RNA-sequencing experiments ^37,80–83^. Data from each study was subjected to re-assignment of cell identity based upon strict criteria (see Methods). **D-F)** UMAP plots showing *LDHA* (D), *LDHB* (E) and *SLC16A1* (F) raw counts clustered according to cell type (data taken from n = 18 donors in ^40,77,78^). **G, H)** Genome browser snapshot of transcription factor binding, histone modification (ChIP-seq, targets as indicated) and RNA-sequencing experiments performed on human islets for LDHA (G) and LDHB (H) ^84^. Scales in A-C represent RPKM.

Recent studies in cancer cells have shown that pyruvate-> lactate conversion is unaffected in single knockouts of LDHA and LDHB, and only in a double LDHA/B knockout is pyruvate no longer converted to lactate ^43,44^. Thus, LDHB can theoretically compensate for LDHA activity when LDHA expression levels are low. However, we cannot exclude the possibility that LDHA may also catalyze pyruvate to lactate conversion in β cells as they do contain detectable LDHA mRNA, based on both single cell RNA-sequencing and bulk RNA-sequencing of FACS purified α cells and β cells (Figure 5A-D). This is consistent with the open chromatin conformation and transcription factor binding to the LDHA promoter in the human islet (Figure 5G). By contrast, *LDHB* possessed two specific enhancers within the CTCF boundaries, suggestive of β cell-specific regulation, although no transcription factor binding was observed at the promoter (Figure 5H).

### LDHB protein localizes predominantly to human **β** cells within the endocrine compartment

To confirm transcriptomic findings, immunohistochemistry was performed using an antibody against LDHB, validated by the human tissue atlas using TMT-MS, protein array and subcellular localization (https://www.proteinatlas.org/ENSG00000111716-LDHB/summary/antibody). To further validate the antibody for use in human β cells, western blotting for LDHB was performed on extracts from EndoC-βH1 cells treated with either siCTL or siLDHB. Confirming antibody specificity, a three-fold reduction in LDHB expression could be seen in siLDHB versus siCTL samples, with knockdown further confirmed by qRT-PCR for mRNA levels (Figure S3A and B).

Similarly to total LDH (i.e. LDHA/LDHB), LDHB was located throughout the cytoplasm as punctate spots (Figure 6A). In line with the transcriptomic analysis and further confirming antibody specificity, LDHB protein could be readily observed throughout the β cell compartment, but was undetectable in the majority (81%) of α cells (Figure 6A-C). We did however notice a small subpopulation (∼19%) of α cells with high levels of LDHB, comparable to those detected β cells (Figure 6B-D). Likewise, a proportion of β cells could be differentiated by their absent or low expression of LDHB (26%) (Figure 6B-D). As for LDH, all results were confirmed with multiple immunostaining runs, using both 40x and 63x objectives (Figure S2A-D).

**Figure 6:**
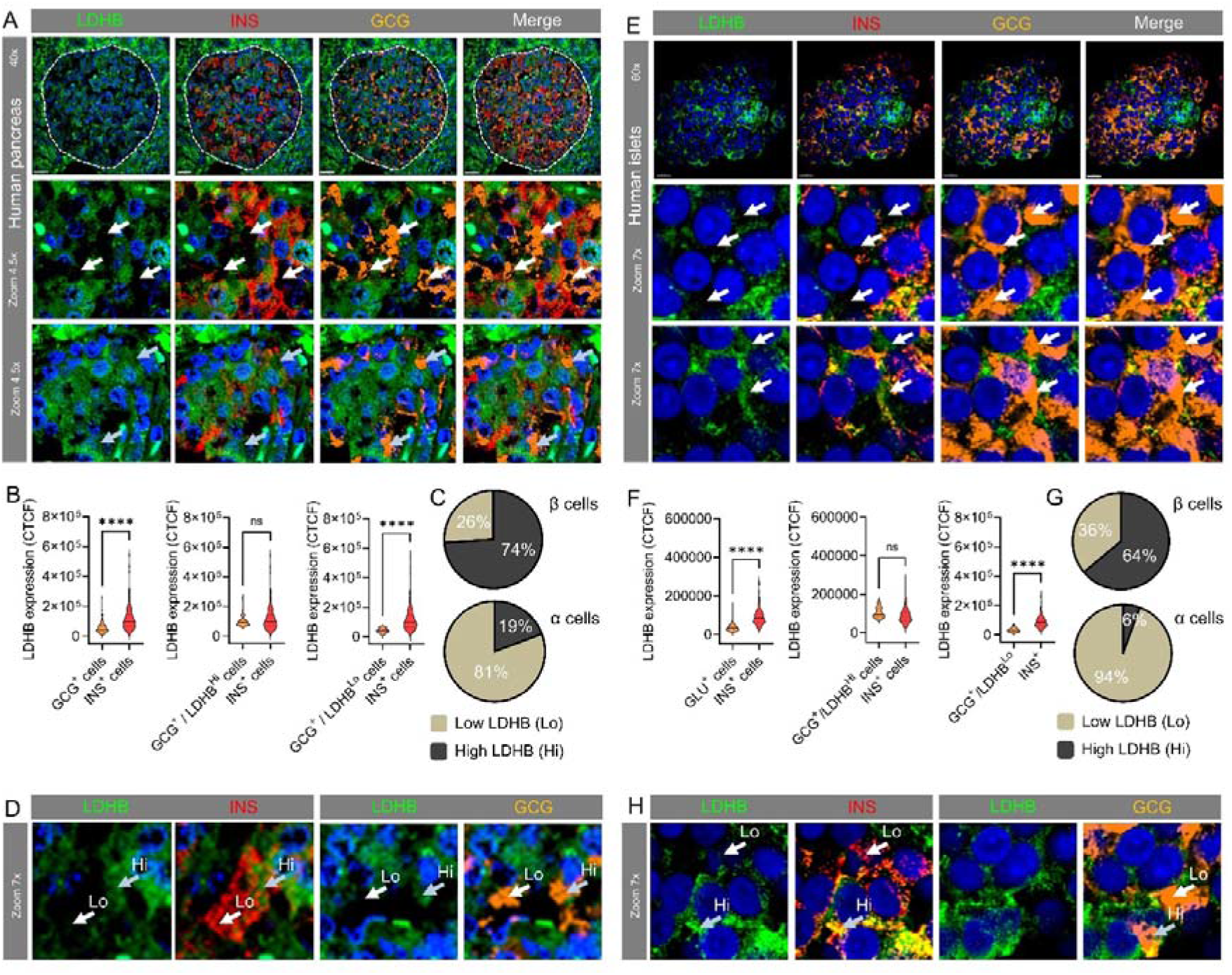
LDHB is predominantly localized to β cells within the islet. **A)** LDHB protein expression strongly co-localizes with insulin (INS) expression in human pancreas sections. Zoom-in (middle panel) shows absence of LDHB in glucagon (GCG)+ cells. Zoom-in (bottom panel) shows a small subpopulation of GCG+ cells with detectable LDHB staining. **B**) Quantification of LDHB expression in GCG+ and INS+ cells in human pancreas sections, including sub-analysis of GCG+ segregated by high (Hi) and low (Lo) LDHB levels (> 8 x 10^4^ CTCF) (n= 150 cells, 3 donors) (Mann-Whitney test). **C**) Pie-charts showing the proportions of GCG+ and INS+ cells that express Hi or Lo LDHB in human pancreas sections (n= 150 cells, 3 donors). **D)** Representative images (zoom-in from A) showing classification of LDHB Hi and Lo cells for GCG and INS. **E)** As for A), but showing that LDHB protein expression remains strongly co-localized with INS in isolated human islets. **F)** Quantification of LDHB expression in GCG+ and INS+ cells in isolated human islets, including sub-analysis of GCG+ segregated by high (Hi) and low (Lo) LDHB levels (> 8 x 10^4^ CTCF) (n= 180 cells, 3 donors) (Mann-Whitney test). **G, H)** As for B and C, but isolated human islets (n= 180 cells, 3 donors). Scale bar = 30 µm. Violin plot shows median and interquartile range.

Since isolated islets were used for metabolic tracing, and pancreas digestion and culture could conceivably influence LDH/LDHB expression levels, immunostaining was repeated in isolated human islets. Almost identical results were obtained in isolated human islets and pancreas sections, suggesting that culture time does not influence LDH/LDHB levels and ergo lactate generation (Figure 6E-H).

### LDHB inhibition slightly increases lactate generation in human **β** cells without affecting the cytosolic ATP/ADP setpoint

While GC-MS- and NMR-based stable isotope tracing provide detailed information about glycolysis and the TCA cycle, this method requires hundreds of islets and is not amenable to purified β cell and α cell fractions.

To establish the contribution of the β cell compartment to lactate generation, we used Ad-RIP-Laconic, a β cell-specific lactate FRET sensor ^45,46^. Experiments in human islets showed that high (17 mM) glucose was able to increase intracellular lactate levels after a lag of ∼ 5 mins (Figure 7A-H). To investigate any role of LDHB, islets were pre-incubated in vehicle or AXKO-0046 for 2 hours, a specific LDHB inhibitor with no detectable activity on LDHA ^47^. AXKO-0046 was used at 10 μM, which corresponds to its maximal inhibitory concentration ^47^. While AXKO-0046 did not appreciably alter glucose-stimulated lactate dynamics, there was a small (10-20%) but replicable increase in glucose-stimulated lactate generation compared to control in islets from all donors examined (Figure 7A-I). Similar results were observed in mouse β cells, which express LDHB albeit at lower levels compared to human β cells (Figure 7J-L).

**Figure 7:**
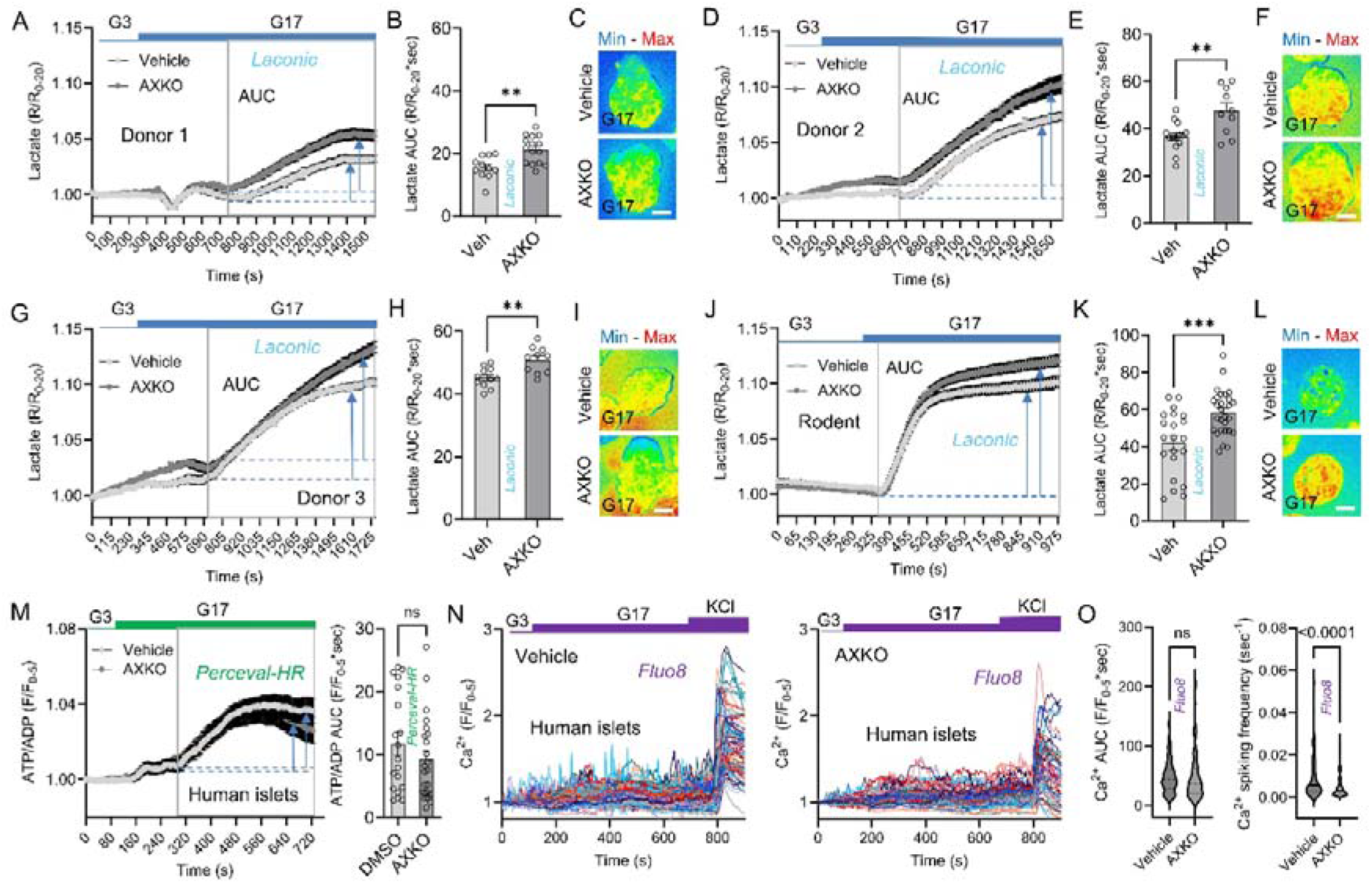
Effects of LDHB inhibition on glucose-stimulate lactate, ATP/ADP and Ca^2+^ in human β cells. **A)** 17 mM glucose stimulates rapid increases in cytosolic lactate in vehicle-treated islets from donor 1, measured using the β cell-specific FRET sensor RIP-Laconic. Glucose-stimulated lactate generation is amplified by an LDHB inhibitor (AXKO-0046) (n = 10-15 islets). **B)** Summary bar graph showing significant amplification and inhibition of lactate by AXKO-0046 (n = 10-15 islets) (one-way ANOVA and Sidak’s post-hoc test). **C)** Representative images showing human β cell lactate levels in the presence of vehicle orAXKO-0046. **D-F)** As for A-C, except donor 2 (n = 10-14 islets) (unpaired t-test). **G-I)** As for A-C, except donor 3 (n = 10-12 islets) (unpaired t-test). **J-L)** AXKO-0046 amplifies glucose-stimulated lactate generation in mouse β cells. **M)** AXKO-0046 does not significantly influence glucose-stimulated ATP/ADP ratios in human β cells, measured using RIP-Perceval-HR (n = 3 donors, 19-23 islets) (unpaired t-test). **N, O)** Individual traces (N) and summary bar graph (O) showing that AXKO-0046 influences Ca^2+-^spiking frequency, but not Ca^2+^ levels in human β cells (n = 3 donors, 87-101 cells) (Mann-Whitney test). Unless other stated, traces shown mean ± SEM. Bar graphs show individual datapoints and mean ± SEM. Scale bar = 83 µm. Arrows and box show AUC calculation boundaries. G3, 3 mM glucose; G17, 17 mM glucose. Islets were pre-incubated with 10 µM AXKO-0046.

Given that LDHB reduces lactate generation, we next looked at whether LDHB inhibition influences glucose-stimulated ATP/ADP and Ca^2+^ rises in human β cells. Vehicle-treated human islets responded to high glucose with increases in ATP/ADP ratios, which were not significantly influenced by LDHB inhibition (Figure 7M). Similarly, glucose-stimulated Ca^2+^ fluxes were identical between vehicle- and AXKO-0046-treated islets (Figure 7N and O). A more detailed analysis of Ca^2+^ oscillations, however, showed a reduction in Ca^2+^ spiking frequency in the presence of AXKO-0046 (Figure 7N and O), in keeping with previous results showing that lactate directly opens K_ATP_ channels ^13^.

Lastly, we attempted to replicate experiments in EndoC-βH5 spheroids, which also express LDHB (Figure S3C). Unexpectedly, EndoC-βH5 spheroids did not tolerate long pre-incubation times with higher concentrations of AXKO-0046, showing signs of blebbing. Thus, spheroids were instead pre-incubated with 100 nM AXKO-0046. At this concentration, AXKO-0046 suppressed both glucose-stimulated ATP/ADP ratios and Ca^2+^ fluxes (Figure S3D-G). Analysis of transcriptomic data revealed that EndoC-βH5 spheroids express moderate LDHA levels, which might explain the exaggerated effects of LDHB inhibition versus primary human β cells (64.11 ± 2.60 versus 247.73 ± 5.00 TPM, *LDHA* versus *LDHB*; mean ± SD; taken from GSE224732 ^48^).

## DISCUSSION

Using ^13^C_6_ glucose labeling coupled to GC-MS and 2D ^1^H-^13^C HSQC NMR spectroscopy, we have been able to obtain a high-resolution map of glucose fate within human and mouse islets. Unexpectedly, both human and mouse islets accumulate m+2 and m+3 lactate, meaning that lactate produced downstream of the TCA cycle (m+2) as well as via pyruvate - > lactate conversion (m+3) contribute equally to overall lactate generation. However, pyruvate-> lactate conversion was much higher (∼ 6-fold) in human compared to mouse islets, in line with relative LDH/LDHB expression levels. Finally, we show that labeled lactate and glutamate accumulate as lactate_110_ and glutamate_00011_, confirming greater flux through PDH versus PC in both species, and the ultimate fate of glucose oxidation in the TCA cycle ^11^. The major findings are schematically represented in Figure 8.

**Figure 8:**
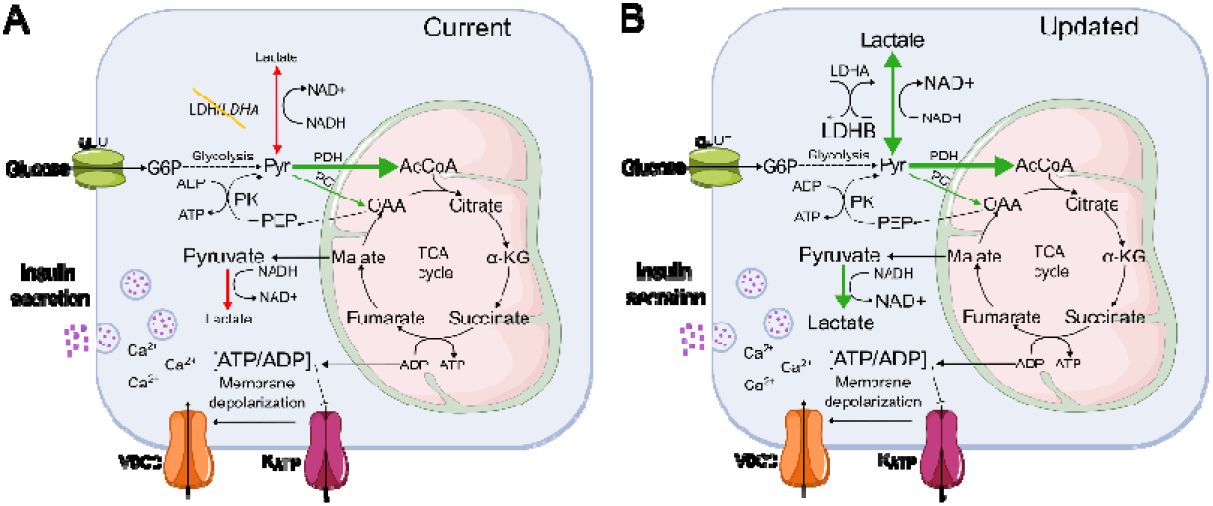
Schematic showing pyruvate management in human islets. **A)** In the current view (top) of human β cell metabolism, glycolytically-derived pyruvate enters the TCA through the actions of pyruvate dehydrogenase (PDH) and pyruvate carboxylase (PC). The PEP cycle and extra-mitochondrial ADP might also contribute to K_ATP_ channel regulation and the triggering phase of insulin secretion. Alternative fates for pyruvate (i.e. generation of lactate) are suppressed due to low levels of LDH/*LDHA* expression. **B)** The high-resolution view of β cell metabolism reveals that LDHB contributes to regulation of lactate generation. The TCA cycle intermediary malate is also converted to pyruvate, which is then further converted to lactate leading to two inter-dependent pools of reducing equivalents to support REDOX (schematic produced in Inkscape, Inkscape Project).

The observation that the islet lactate pool is derived from both TCA cycle- and pyruvate-derived sources suggests that mechanisms must be in place for direct pyruvate conversion. In many tissues, pyruvate would be converted to lactate by LDH, however, lactate generation is minimal in purified rat beta cells ^49^, and the *Ldha* isozyme of the enzyme has been shown to be disallowed or absent in the murine pancreatic β cell ^8,10,50^. In keeping with these findings and further supporting a more limited role for LDH in rodents, lactate and LDH protein could only be detected at low levels in mouse islets, although glucose-stimulated lactate rises could still be detected by us and others in single β cells^46^. By contrast, LDH expression in human β cells was ∼2-fold higher than in mouse β cells, albeit less than that in the human liver, known to strongly express LDH. Analysis of multiple published scRNA-seq datasets further showed that human β cells specifically and strongly express *LDHB* within the islet, which encodes the beta subunit of LDH, confirming previous single cell screening studies by van Gurp et al ^40^. LDHA expression was also detected at low-moderate levels based upon bulk RNA-seq of purified human β cells.

Oxygen availability and ergo NADH:NAD^+^ are both known to influence affinity of LDHA and LDHB for their substrates, with anaerobic conditions favoring pyruvate to lactate conversion and vice versa under normoxia. Notably, glucose-stimulation leads to large increases in beta cell O_2_ consumption sufficient to activate HIF enzymes ^51,52^. Supporting the findings here, recent studies using Seahorse assays have shown that glucose-stimulated oxygen consumption ratio is ∼ 30% higher in mouse versus human islets due to higher leak (with caveats about inter-islet heterogeneity noted) ^53^. As such, for a given O_2_ supply, mouse islets consume more O_2_, which should lead to increased lactate generation. However, lactate generation remains low compared to human islets, since mouse islets express less LDHA/LDHB.

It is possible that α-cells contribute to the accumulation of lactate. Human α cells account for ∼35% of the entire islet and express *LDHA* at levels six times higher than β cells ^54,55^. However, a major source of α cell lactate is via monocarboxylate transporters ^9,56,57^, which are unlikely to play a role here as lactate was absent from the tracing medium. In addition, while the total amount of lactate was only doubled in humans compared to mice, the m+3 (TCA-derived) and m+3 (glycolytically-derived) lactate accumulation was ∼six-fold higher in human *versus* mouse islets, which cannot be accounted for solely by differences in α cell proportion. Total LDH levels were found to be similar in human α-cells and β cells, although a recent study suggests that plasma membrane-compartmentalized LDH activity differs between human α and β-cells, and plays a more significant role in α-cells ^13^. Lastly, studies with Laconic, a lactate FRET sensor, showed that human β cells are capable of generating significant intracellular lactate levels in response to stimulation with high glucose concentration. Mouse β-cells were also found to generate cytosolic lactate in response to glucose (also ^46^) using the same approach, but the species were not quantitatively compared. Taken together, these data suggest that α cell lactate only makes a very minor contribution to the whole m+3 and m+2 lactate increase detected here. We note that human islet preps also typically comprise ∼ 50% acinar, duct and other non-islet cells, where LDHA/LDHB is enriched versus islet cells (see Figure 3D and E). While islets were hand-picked, filtered and cultured for 2-3 days during which acinar cells are expected to die off, we cannot exclude a contribution of the exocrine and other compartments to the findings here.

Human and mouse islets display a greater accumulation of lactate_110_, rather than lactate_011_. While the accumulation of lactate_110_ is indistinguishable in the PDH- and PC-mediated TCA cycle, the 011 isotopomer would only derive from the reductive metabolism of PC-derived glutamate. Corroborating this, in islets from both sschuiecies, the major glutamate isotopomer derived from exogenous ^13^C_6_ glucose was glutamate_00011_. Although glutamate is not a TCA cycle metabolite, it is in rapid exchange with α-KG and can be used as a read-out of TCA cycle flux through PDH or PC. Consequently, the accumulation of glutamate_00011_ provides further evidence for a higher reliance of the TCA cycle on the activity of PDH, rather than PC. Although PC and PDH were thought to contribute equally to the TCA cycle in β-cells ^20–22^, previous studies have shown that high glucose concentrations in vitro, more reflective of those seen post-prandially in vivo, are associated with an increase toward PDH activity ^11,15,34^. Our studies thus show that the relative contribution of PC to the TCA cycle is much lower than PDH (∼20%), confirming findings from Alves et al in glucose-traced INS-1 cells ^15^ and Lewandowski et al in human islets ^11^. This does not mean that PC flux is unimportant, since PC is much more responsive to glucose stimulation than PDH ^11^.

While anaplerosis through PC is relatively limited in the β cell, we note that glucose carbons can repeatedly transit the PEP cycle to increase ATP/ADP in the cytosol ^11^ (before pyruvate is ultimately oxidized in the TCA cycle). As such, PC is able to make disproportionate contributions to K_ATP_ channel closure, and hence the triggering phase of insulin secretion, by generating plasma-membrane localized increases in ATP/ADP ^14^. Moreover, PC might make relatively more important contributions to anaplerosis in stressed human beta cells in which increased PC activity reduces NO production to counteract inflammation ^58^. PC is also important for funneling pyruvate into generation of the GSIS-coupling factors citrate and isocitrate, which sustain 2-ketoglutrate and NADPH production ^18^. Nonetheless, insulin secretion is not the only energy sink on the β cell, and glucose flux through PDH is likely to provide a source of acetyl-CoA to support other demands such as continued insulin release and protein synthesis ^49^. Indeed, insulin biosynthesis and ion pumping have both been shown to be greater β cell energy sinks than exocytosis ^59^.

What might be the role of direct pyruvate to lactate conversion in pancreatic β cells? Since the action of LDH leads to oxidation of NADH to NAD^+^, lactate accumulation could provide a source of reducing equivalents to support other NADH-producing metabolic pathways. Providing evidence for an important contribution of pyruvate to lactate conversion in NADH/NAD+ balance, the most abundant lactate isotopomers were lactate_111_ and lactate_110_, which respectively represent lactate generation upstream and downstream of the TCA cycle. Since the conversion of pyruvate-> lactate is associated with the generation of cytosolic NAD^+^, higher levels of total lactate_111_ in humans might reflect an increase in the activity of NADH-producing pathways relative to rodents. Notably, the alanine isotopomer distribution showed accumulation of alanine_000_ and alanine_111_, suggesting that pyruvate accumulated downstream of the TCA cycle is employed in the regeneration of lactate_110_. Arguing against this possibility, β cells lack the capacity to extracellularly transport lactate, and are thought to already have a large capacity to produce reducing equivalents, e.g. via the glycerol phosphate and malate-aspartate shuttles ^60^. Although membrane-associated LDH was found to form nanodomains with K_ATP_ channels and support the local production of NAD^+^ for GADPH, the contribution of LDH activity (at least in mouse β-cells) was small relative to other unidentified plasma membrane-associated NADH oxidases that support this function. Lastly, significant pools of cytosolic pyruvate and lactate, and sufficient LDH to equilibrate these pools, might provide a buffer to minimise wide fluctuations in NADH/NAD (or NAD(P)H/NADP), offering some protection against oxidative stress ^61^.

In summary, by combining MID and multiplet analyses, we show that glucose makes a similar contribution to glycolysis and the TCA cycle in human and mouse islets. Furthermore, the isotopomer distribution confirms that, in both species, the relative activity of PDH is higher than that of PC at elevated glucose concentration. However, the generation of fully-labeled lactate was found to be much higher in human versus mouse islets, with LDHB and acting to limit lactate levels. Together, these results demonstrate that lactate generation should be reconsidered in light of beta cell development, metabolism and function.

### Limitations

There are a number of limitations in the present studies. Firstly, glucose-tracing studies in purified α cells and β cells are warranted, although should be interpreted in light of loss of cell-cell interactions, changes in cell phenotype, as well as input material required. Nonetheless, our data provide a detailed and interrogable map of glucose metabolism pertaining to steady state insulin release in human and mouse islets, as well as other critical β cell housekeeping functions. Secondly, glucotoxicity might induce the upregulation of disallowed genes in β cells ^62^. However, it is unlikely that the timings (12 hrs) and glucose concentration (10 mM) used here would overtly influence human β cell lactate generation, since *LDHA* or *LDHB* were not found to be differentially expressed in human islets exposed to 22.2 mM glucose for 4 days ^63^. Thirdly, to increase resolution of glucose fate-mapping, tracing should be performed at different timepoints, similarly to recent studies ^32,33^. Such studies are at the moment limited by 800 MHz NMR access and Helium availability. Fourthly, functional studies in primary human beta cells deleted for LDHA/LDHB are warranted, although there are concerns that the other LDH isoenzymes might be upregulated over chronic timecourses. Lastly, the effects of LDHB inhibition on glucose-stimulated lactate generation were relatively small. This could reflect the potency of AXKO-for LDHB in the tissue setting, existence of lactate microdomains, or the high glucose concentration used. Future studies looking at glucose concentration curves, as well as tracing-based analyses will be valuable.

## STAR METHODS

## RESOURCE AVAILABILITY

### Lead contact

Further information and requests for resources and reagents should be directed to and will be fulfilled by the lead contact, David J. Hodson.

### Materials availability

This study did not generate any new unique reagents.

### Data and code availability

This paper analyses existing, publicly available data. Accession numbers for the datasets are listed in the key resources table.

All data reported in this paper will be shared by the lead contact upon request.

## EXPERIMENTAL MODEL AND STUDY PARTICIPANT DETAILS

### Mice

Male 8 to 12 week-old CD1 mice (Charles River stock no. 022) were used as tissue donors. Mice were socially-housed in specific-pathogen free conditions under a 12 hour light-dark cycle with ad libitum access to food and water, relative humidity 55 ± 10% and temperature 21 ± 2 °C.

Animal studies were regulated by the Animals (Scientific Procedures) Act 1986 of the U.K. (Personal Project Licences P2ABC3A83 and PP1778740). Approval was granted by the University of Birmingham and University of Oxford Animal Welfare and Ethical Review Bodies (AWERB).

### Human

Human islets (Lille): human pancreatic tissues were harvested from brain-dead adult donors in accordance with the Lille clinical islet transplantation program’s traceability requirements (clinicaltrials.gov, NCT01123187, NCT00446264, NCT01148680), and were approved in agreement with French regulations and the Ethical Committees of the University of Lille and the Centre Hospitalier Régional Universitaire de Lille.

Human islets (Milan): the use of human islets for research was approved by the Ethics Committee of San Raffaele Hospital in Milan (IPF002-2014).

Human islets (Oxford): human pancreata were retrieved from Donors after Brain-Stem Death (DBD) with appropriate consent and ethical approval under 09/H0605/2 (REC: Oxfordshire Rec B). Islets were isolated in the DRWF Oxford Human Islet Isolation Facility using established isolation methods.

Human pancreas sections (Oxford): post-mortem pancreas samples were obtained from donors, with appropriate permissions registered under CUREC R83564/RE001.

Studies with human islets and pancreata were approved by the University of Birmingham Ethics Committee, the University of Oxford Ethics Committee, as well as the National Research Ethics Committee (REC 16/NE/0107, Newcastle and North Tyneside, UK).

Liver sections (Oxford): Authorisation for research use of steatotic livers unsuitable for transplantation was obtained by a specialist nurse in organ donation in accordance with NHSBT guidelines, with appropriate permissions registered under REC 14/LO/0182. The study was approved by the North East – Tyne and Weir South research ethics committee (16/NE/0248) and by the NHSBT Research, Innovation and Novel Technologies Advisory Group.

## METHOD DETAILS

### Study design

No data were excluded, and all individual data points are reported in the figures. The measurement unit is animal or donor, with experiments replicated independently on different days. Islet isolation is a nuisance variable and as such data are taken from independent islet preparations. Samples and animals were allocated to treatment groups in a randomized manner to ensure that all states were represented in the different experiment arms. MID analysis was performed by a user blinded to sample identity. For metabolic tracing, nine samples per investigated state are required to correctly reject the null hypothesis (two-tailed test) for an effect size (d) = 1.5 (calculated in GPower 3.1).

### Mouse islets

Animals were culled using a schedule-1 method followed by injection of the common bile duct with 1 mg/mL collagenase NB 8 (Serva) in RPMI 1640 (Gibco) and pancreas dissection. After dissection, the pancreas was incubated in a water bath at 37°C for 12 min. Subsequently, the tissues were shaken in 15 mL of RPMI 1640 and centrifuged for 1 min at 1500 rpm three times to induce mechanical digestion. Islets were separated using Histopaque-1119 and 1083 (Sigma-Aldrich) gradients, before hand-picking and culture. Unless otherwise stated, the islets obtained were kept in culture in RPMI 1640 supplemented with 10% fetal bovine serum (FBS, Gibco), 100 units/mL penicillin, and 100 μg/mL streptomycin (Sigma-Aldrich), at 37°C and 5% CO_2_.

For immunohistochemistry, sections were cut from formalin-fixed, paraffin-embedded (FFPE) pancreata obtained from wild-type male and female C57BL6 mice fed standard diet or high fat diet (60% fat, Research Diets, catalog D12492) for 8-12 weeks. To reduce the number of animals used in experiments in line with NC3Rs policy, blocks were re-used from a previous study ^64^, new sections cut at different depth, and immunostained with different markers (LDH, PDX1, see below).

### EndoC-**β**H1 and EndoC-**β**H5 cells

Total protein was extracted from provided EndoC-βH1 cells using RIPA lysis buffer. Lysis buffer was supplemented with EDTA-free protease inhibitor and phosphatase inhibitor cocktails. Lysates were resolved on SDS–PAGE, transferred to a nitrocellulose membrane using the Trans-Blot Turbo Transfer Pack, blocked in 5% milk for 1 hr, followed by an incubation with a polyclonal Anti-LDHB antibody (HPA019007) at 4 °C overnight. After washing, membrane was then incubated for 2 hr at room temperature with secondary antibody conjugated to horseradish peroxidase (HRP). After incubation with Clarity Western ECL Substrate, bands were detected with Bio-Rad GelDoc Biosystems Western Blotting imager.

EndoC-βH1 cells were transfected using Lipofectamine RNAiMAX with ON-TARGETplus siRNA targeting *LDHB* (L-009779-00, Dharmacon, Horizon Discovery) at a final concentration of 80nM. Non-targeting control pool (D-001810-10, Dharmacon, Horizon Discovery) was used as a control. Briefly, siRNA and Lipofectamine RNAiMAX were combined in OptiMEM and applied to the cells. Medium was changed 2.5h later for fresh EndoC-βH1 culture medium. Cells were harvested in RIPA lysis buffer or in RLT lysis buffer for mRNA purification (Qiagen).

Lysis buffer was supplemented with EDTA-free protease inhibitor and phosphatase inhibitor cocktails. Lysates were resolved on SDS–PAGE, transferred to a nitrocellulose membrane using the Trans-Blot Turbo Transfer Pack, blocked in 5% milk for 1 hr, followed by an incubation with a polyclonal Anti-LDHB antibody (HPA019007) at 4 °C overnight. After washing, membranes were then incubated for 2 hr at room temperature with secondary antibody conjugated to horseradish peroxidase (HRP). After incubation with Clarity Western ECL Substrate, bands were detected with Bio-Rad GelDoc Biosystems Western Blotting imager.

Primers used for PCR were as follows: PPIA (housekeeping) Fw primer: ATGGCAAATGCTGGACCCAACA, PPIA (housekeeping) Rv primer: ACATGCTTGCCATCCAACCACT, LDHB Fw primer: TGATGGATCTGCAGCATGGG and LDHB Rv primer: CAGATTGAGCCGACTCTCCC.

To generate EndoC-βH5 (Human Cell Design) pseudoislets, 1 million cells were seeded in Aggrewell Multiwell plates in ULT1β1 culture media (Human Cell Design) at 37°C and 5% CO2. Pseudoislets aggregated in 3-4 days and media was changed every 2 days.

### Human islets

Islets were cleared of possible debris via filtration with a 40 μm cut-off filter, hand-picked and cultured in CMRL medium (Corning) containing 5.5 mM glucose, 10% FBS, 100 units/mL penicillin, 100 μg/mL streptomycin and 0.1% amphotericin B (Sigma-Aldrich) at 37°C and 5% CO_2_. Human donor characteristics are reported in Table S1.

### ^13^C_6_ glucose tracing

For ^13^C_6_ glucose tracing, 60 (for GC-MS) or 150-230 (for NMR) islets were used. Isolated islets were cultured in RPMI 1640, no glucose medium (Gibco), supplemented with 10% BSA, 10% FBS, 100 units/mL penicillin, and 100 μg/mL streptomycin plus 10 mM ^13^C_6_ glucose (Sigma-Aldrich). After 24 h, the metabolites were extracted adding HPLC-grade methanol, HPLC-grade distilled H_2_O containing 1 μg/mL D6-glutaric acid and HPLC-grade chloroform (all from Sigma-Aldrich) in a 1:1:1 ratio, to the islets. Following centrifugation, the polar fractions were collected and vacuum dried before either GC-MS or NMR analyses. A 24 h tracing duration was used to allow steady state labelling of both glycolytic and TCA metabolites in the same sample, as well as sufficient ^13^C incorporation for NMR-based labelling pattern annotation. While antibiotics have been shown to influence mitochondrial function, research islet isolation is a non-sterile procedure. Antibiotic use is therefore justified, since low grade infection is likely to exert a larger (and unnoticed) effect on mitochondrial activity.

### GC-MS

The dried polar extracts were prepared for GC-MS analysis through solubilization in 40 µL of 2% methoxyamine hydrochloric acid in pyridine (Fisher Scientific) at 60°C for 60 min and derivatization with 60 µL of N-tertbutyldimethylsilyl-N-methyltrifluoroacetamide (MTBSTFA) with 1% tertbutyldimethyl-chlorosilane (TBDMCS) (both from Sigma-Aldrich). The suspension was further incubated at 60°C for 60 min, before being centrifuged at 13300 rpm for 10 min at 4°C and transferred to chromatography vials with a glass insert (Restek) for GC-MS analysis. The samples were analyzed on an Agilent 8890 GC and 5977B MSD system. To do this, 1 μL of sample was injected in splitless mode with helium carrier gas at a rate of 1.0 mL/min. The compound detection was carried out in scan mode and total ion counts of each metabolite were normalized to the internal standard D6-glutaric acid and corrected for natural ^13^C abundance ^65^. Retention times, ion counts and MID data are provided in Table S2 (mouse) and Table S3 (human).

### NMR spectroscopy

Following the ^13^C_6_ glucose tracing, the dried polar metabolites were resuspended in 60 µL of phosphate buffer: 57.8 mM disodium phosphate (Na_2_HPO_4_, Sigma-Aldrich), 42.2 mM monosodium phosphate (NaH_2_PO_4_, Sigma-Aldrich), 0.5 mM 3-(trimethylsilyl) propionic-2,2,3,3-d4 acid sodium salt (D4-TMSP, Sigma-Aldrich) in deuterium oxide (D_2_O, Sigma-Aldrich). Subsequently, the samples were centrifuged for 10 min at 14800 rpm and sonicated in an ultrasonic bath for 5 min, before being loaded into NMR tubes (outer diameter: 1.7 mm, Bruker) for acquisition. A Bruker Neo 800 MHz NMR spectrometer equipped with a 1.7 mm z-PFG TCI Cryoprobe was used to acquire 2D ^1^H,^13^C-HSQC NMR spectra. The HSQC spectra were acquired with echo/anti-echo gradient coherence selection with an additional pre-saturation for suppressing the residual water resonance. The spectral widths were 15.6298 ppm and 189.7832 ppm in the ^1^H and ^13^C dimension, 512 complex data points were acquired for the ^1^H dimension and 25% (512) out of 2048 complex data points were acquired for the ^13^C indirect dimension using a non-uniform sampling scheme. Apparent ^13^C,^13^C J-coupling was enhanced four-fold. The interscan relaxation delay was set to 1.5 s. 2D ^1^H,^13^C-HSQC spectra were reconstructed via the compressed sensing IRLS algorithm using the MddNMR (version 2.5) ^66^ and NMRPipe (version 9.2) ^67^ software. All NMR spectra were analyzed in the MATLAB based MetaboLab software package ^68^.

### Immunohistochemistry

Formalin-fixed, paraffin-embedded pancreas, liver and isolated islets were cut at 5µm and deparaffinised. Human donor characteristics are reported in Table S4. PBS with 2% BSA and 0.2% Triton X was used to block sections for 1 hour at room temperature. Heat induced antigen retrieval was performed with 10 mM citrate buffer (pH6) using a microwave for 10 minutes and cooled for 10 minutes in cold water. Sections were incubated with primary antibodies overnight at room temperature, followed by washes with PBS-2% BSA and 0.2% Triton X. Secondary antibodies were incubated for 2 hours at room temperature. Sections were mounted with DAPI (Vectashield, cat no. H-1500) and stored at 4°C. Primary antibodies used were mouse anti-LDH (Santa Cruz Biotechnology, cat no. sc-133123, RRID: AB_2134964), rabbit polyclonal anti-LDHB (Sigma-Aldrich, cat no. HPA019007, RRID:AB_2670008), guinea-pig anti-PDX1 (Abcam, cat# ab47308, RRID:AB_777178), guinea pig anti-insulin (Abcam, cat no. ab7842, RRID: AB_306130), and mouse monoclonal anti-glucagon (Sigma-Aldrich, cat# G2654, RRID:AB_259852). Human and mouse LDHA, LDHB and LDHC share 94.0%, 97.9% and 74.5% protein sequence identity, respectively. We note however that LDHC is undetectable in the human pancreas ^35^ and is largely confined to the testis ^36^. For LDHA and LDHB detection, we used an amplification step consisting of anti-mouse biotinylated (DAKO, cat no. E0413) or anti-rabbit biotinylated (NOVEX, cat no. A16039, RRID: AB_2534713) antibodies, followed by incubation with streptavidin-FITC (Sigma-Aldrich, Cat# S-3762). For all other markers, secondary antibodies used were anti-guinea pig Alexa Flour 568 (Thermo Fisher, cat no. A-11075, RRID: AB_141954), anti-guinea pig Alexa Flour 647 (Thermo Fisher, cat no. A-21450, RRID: AB_141882) and anti-mouse Alexa Flour 568 (Thermo Fisher, cat no. A-11004, RRID: AB_2534072).

Sections were imaged using an Olympus FV3000 confocal microscope equipped with high-sensitivity spectral detectors and 40x, 1.25 NA and 60x, 1.30 NA silicone objectives. Excitation and emission wavelengths for DAPI, Alexa Fluor 488, Alexa Flour 568 and Alexa Flour 647 were λex = 405 nm / λem = 406 – 461 nm, λex = 488 nm / λem = 499 – 520 nm, λex = 561 nm / λem = 579 – 603 nm, λex = 640 nm / λem = 649 - 700 nm respectively.

### Lactate, Ca^2+^ and ATP/ADP imaging

Isolated islets were transduced 24-48 hrs with the beta cell specific lactate FRET sensor, Ad-RIP-Laconic ^45,46^, prior to widefield imaging using a Nikon Ti-E base equipped with 89 North LDI-7 Laser Diode Illuminator, 25x / 0.8 NA objective and Prime BSI Express sCMOS. Excitation was delivered at λ===445=nm and emission detected at λ===460–500 and λ===520–550=nm for mTFP and Venus, respectively. All experiments were performed in HEPES-bicarbonate buffer containing (in mmol/L) 120 NaCl, 4.8 KCl, 24 NaHCO3, 0.5 Na2HPO4, 5 HEPES, 2.5 CaCl2, 1.2 MgCl2, supplemented with 3–17 mmol/L d-glucose, and bubbled with 95% O_2_ / 5% CO_2_. Laconic intensity was calculated as the ratio of mTFP/Venus. AXKO-0046 (MedChemExpress, cat no. HY-147216) was used as a selective nanomolar-affinity small molecule inhibitor of LDHB (IC_50_ = 42 nM, IC_MAX_ = 10^-5^ nM).

Ca^2+^ and ATP/ADP imaging was performed in human islets and EndoC-βH5 spheroids loaded with Fluo8 or transduced with Ad-RIP-Perceval-HR, respectively ^11,69^. Islets were imaged as above, with excitation delivered at λ====470 nm using a CrestOptics X-light V2 spinning disk head; and emission detected at λ===500-550.

### Image analysis

Images were deconvolved using Imaris Clearview (Oxford Instruments) and an AMD Radeon Pro W5500 GPU with 8GB GDDR6. Image analysis was performed in ImageJ (NIH) using corrected total cell fluorescence (CTCF), which is the integrated density corrected for the product of cell area and mean background fluorescence ^70,71^.

### Transcriptomics analysis

Quantification data for all published scRNA-seq datasets were kindly provided by Leon Van Gurp ^40^. In brief, pseudo-counts were normalized for each data set using Seurat ^72^, and the cell identity was assigned based on the requirement for hormone gene expression to be in the top 1% expressed genes in each cell using Aucell ^73^. Quantification for published FACS sorted alpha and β cells were obtained from GEO database repository under Arda et al ^74^. The raw read files for each cell type were merged, trimmed and the transcripts were quantified using Kallisto ^75^ or aligned and quantified as previously described ^76^, with similar results.

For Uniform Manifold Approximation Projection (UMAP) plots, datasets were downloaded from GSE150724 ^40^, GSE84133 ^77^ and GSE114297 ^78^. GSE84133 ^77^ contained a single donor with type 2 diabetes, excluded here. Seurat R package was used to filter, cluster, and identify cell types. Cells were filtered if they had less than 500 unique genes and if more than 20% of their UMI counts were of mitochondrial origin. Only moderate filtering of mitochondrial DNA was conducted because of the nature of the cells. After filtering, raw UMI counts were normalized by scaling to 10,000 followed by log2-transformation. Normalized gene expression was used to find the top 2000 most variable genes. These genes were ranked based on the number of samples they were deemed a top variable gene, scaled to generate z-scores and used to run Principal component analysis (PCA). The first 50 principal components were used to define anchors between datasets using Reciprocal PCA (RPCA). RPCA was run with Donor 1 from ^40^ as a reference. Subsequently, PCA was run on the integrated data and a neighborhood graph built using 20 principal components. Clustering was carried out using a resolution of 0.2 and visualized in 2D space (Uniform Manifold Approximation Plot) for cluster identification. Cells clustered according to their cell type, as marked by their expression of INS, GCG and SST. Most cells in the original clusters had ambient levels of the non-identifying hormone, which might reflect cross-contamination during FACs ^79^. However, there were cells with low expression of the cluster-identifying hormone, and cells with high expression of the non-identifying hormones. These cells were excluded on the premise that they represent doublets, or misclustered cells. After cell type identification, raw counts from each cell were aggregated (summed) according to cell type and donor for ‘pseudobulk’ representation. Markers used for cell type identification are listed in Table S5.

## QUANTIFICATION AND STATISTICAL ANALYSIS

GraphPad Prism 9 (version 9.2.0) was used for all statistical analyses. Data distribution was assessed using the D’Agostino-Pearson normality test. Pairwise comparisons were made using Welch’s test (assuming non-equal standard deviation between groups). Multiple interactions were determined using one-way ANOVA or two-way ANOVA, with Sidak’s post-hoc test. For non-parametric data, pairwise comparisons were made using Mann-Whitney test, and multiple interactions determined using Kruskal-Wallis test, with Dunn’s post-hoc test. Individual datapoints are shown in bar graphs. Unless otherwise stated in the figure legend, all error bars represent mean ± S.E.M. and a p-value less than 0.05 was considered significant: *p< 0.05; **p< 0.01; ***p< 0.001, ****p<0.0001.

## AUTHOR CONTRIBUTIONS

F.C., K.V., A.S.H. performed experiments, analyzed data and wrote the manuscript. D.N., A.C., C.F-M. and H.R.S. performed experiments. Z.J. and I.A. performed bioinformatic analysis. R.N., L.P., C.B., F.P., J.K-C., P.R.V.J. and R.S. isolated and provided human islets. C.L. performed ^1^H,^13^C-HSQC NMR experiments and analysis. G.G.L., D.T., A.B., B.M. and J.R. ran GC-MS on ^13^C_6_ glucose labelled samples and provided analysis. L.H., A.C., C.D.L.C. provided human liver samples. A.P.D. and G.A.R. provided scRNA-seq analysis. R.S. and M.O. participated in antibody validation. M.J.M. provided Ad-RIP-Laconic and also advised on lactate imaging studies. D.J.H. supervised the studies, provided analysis and wrote the manuscript with input from all authors. All authors read and approved the studies.

## ACKNOWLEDGEMENTS

The authors thank Prof. Patrick E. MacDonald and Dr. Jocelyn Manning Fox (Alberta Diabetes Institute IsletCore at the University of Alberta in Edmonton) for provision of human pancreas sections. We would like to acknowledge the support and resources of the Birmingham Metabolic Tracer Analysis Core.

## REFERENCES

1. De Vos, A., Heimberg, H., Quartier, E., Huypens, P., Bouwens, L., Pipeleers, D., and Schuit, F. (1995). Human and rat beta cells differ in glucose transporter but not in glucokinase gene expression. Journal of Clinical Investigation 96, 2489–2495. 10.1172/jci118308.

2. Thorens, B., Sarkar, H.K., Kaback, H.R., and Lodish, H.F. (1988). Cloning and functional expression in bacteria of a novel glucose transporter present in liver, intestine, kidney, and beta-pancreatic islet cells. Cell 55, 281–290. 10.1016/0092-8674(88)90051-7.

3. Rorsman, P., and Ashcroft, F.M. (2018). Pancreatic beta-Cell Electrical Activity and Insulin Secretion: Of Mice and Men. Physiol Rev 98, 117–214. 10.1152/physrev.00008.2017.

4. Rutter, G.A., Pullen, T.J., Hodson, D.J., and Martinez-Sanchez, A. (2015). Pancreatic beta-cell identity, glucose sensing and the control of insulin secretion. Biochem J 466, 203–218. 10.1042/BJ20141384.

5. Henquin, J.C. (2000). Triggering and amplifying pathways of regulation of insulin secretion by glucose. Diabetes 49, 1751–1760.

6. Ferdaoussi, M., Dai, X., Jensen, M.V., Wang, R., Peterson, B.S., Huang, C., Ilkayeva, O., Smith, N., Miller, N., Hajmrle, C., et al. (2015). Isocitrate-to-SENP1 signaling amplifies insulin secretion and rescues dysfunctional beta cells. J Clin Invest 125, 3847–3860. 10.1172/JCI82498.

7. Ainscow, E.K., Zhao, C., and Rutter, G.A. (2000). Acute overexpression of lactate dehydrogenase-A perturbs beta-cell mitochondrial metabolism and insulin secretion. Diabetes 49, 1149–1155. DOI 10.2337/diabetes.49.7.1149.

8. Sekine, N., Cirulli, V., Regazzi, R., Brown, L.J., Gine, E., Tamarit-Rodriguez, J., Girotti, M., Marie, S., MacDonald, M.J., Wollheim, C.B., and, et al. (1994). Low lactate dehydrogenase and high mitochondrial glycerol phosphate dehydrogenase in pancreatic beta-cells. Potential role in nutrient sensing. Journal of Biological Chemistry 269, 4895–4902.

9. Schuit, F., Van Lommel, L., Granvik, M., Goyvaerts, L., de Faudeur, G., Schraenen, A., and Lemaire, K. (2012). beta-cell-specific gene repression: a mechanism to protect against inappropriate or maladjusted insulin secretion? Diabetes 61, 969–975. 10.2337/db11-1564.

10. Pullen, T.J., Khan, A.M., Barton, G., Butcher, S.A., Sun, G., and Rutter, G.A. (2010). Identification of genes selectively disallowed in the pancreatic islet. Islets 2, 89–95. 10.4161/isl.2.2.11025.

11. Lewandowski, S.L., Cardone, R.L., Foster, H.R., Ho, T., Potapenko, E., Poudel, C., VanDeusen, H.R., Sdao, S.M., Alves, T.C., Zhao, X., et al. (2020). Pyruvate Kinase Controls Signal Strength in the Insulin Secretory Pathway. Cell Metabolism 32, 736–750.e735. 10.1016/j.cmet.2020.10.007.

12. Foster, H.R., Ho, T., Potapenko, E., Sdao, S.M., Huang, S.M., Lewandowski, S.L., VanDeusen, H.R., Davidson, S.M., Cardone, R.L., Prentki, M., et al. (2022). β-cell deletion of the PKm1 and PKm2 isoforms of pyruvate kinase in mice reveals their essential role as nutrient sensors for the KATP channel. eLife 11. 10.7554/eLife.79422.

13. Ho, T., Potapenko, E., Davis, D.B., and Merrins, M.J. (2023). A plasma membrane-associated glycolytic metabolon is functionally coupled to K(ATP) channels in pancreatic alpha and beta cells from humans and mice. Cell Rep 42, 112394. 10.1016/j.celrep.2023.112394.

14. Merrins, M.J., Corkey, B.E., Kibbey, R.G., and Prentki, M. (2022). Metabolic cycles and signals for insulin secretion. Cell Metabolism 34, 947–968. 10.1016/j.cmet.2022.06.003.

15. Alves, T.C., Pongratz, R.L., Zhao, X., Yarborough, O., Sereda, S., Shirihai, O., Cline, G.W., Mason, G., and Kibbey, R.G. (2015). Integrated, Step-Wise, Mass-Isotopomeric Flux Analysis of the TCA Cycle. Cell Metab 22, 936–947. 10.1016/j.cmet.2015.08.021.

16. Farfari, S., Schulz, V., Corkey, B., and Prentki, M. (2000). Glucose-regulated anaplerosis and cataplerosis in pancreatic beta-cells: possible implication of a pyruvate/citrate shuttle in insulin secretion. Diabetes 49, 718–726.

17. Lu, D., Mulder, H., Zhao, P., Burgess, S.C., Jensen, M.V., Kamzolova, S., Newgard, C.B., and Sherry, A.D. (2002). 13C NMR isotopomer analysis reveals a connection between pyruvate cycling and glucose-stimulated insulin secretion (GSIS). Proc Natl Acad Sci U S A 99, 2708–2713. 10.1073/pnas.052005699.

18. Joseph, J.W., Jensen, M.V., Ilkayeva, O., Palmieri, F., Alárcon, C., Rhodes, C.J., and Newgard, C.B. (2006). The Mitochondrial Citrate/Isocitrate Carrier Plays a Regulatory Role in Glucose-stimulated Insulin Secretion. Journal of Biological Chemistry 281, 35624–35632. 10.1074/jbc.M602606200.

19. Zhang, G.-F., Jensen, M.V., Gray, S.M., El, K., Wang, Y., Lu, D., Becker, T.C., Campbell, J.E., and Newgard, C.B. (2021). Reductive TCA cycle metabolism fuels glutamine-and glucose-stimulated insulin secretion. Cell Metabolism 33, 804–817.e805. 10.1016/j.cmet.2020.11.020.

20. Cline, G.W., Lepine, R.L., Papas, K.K., Kibbey, R.G., and Shulman, G.I. (2004). 13C NMR isotopomer analysis of anaplerotic pathways in INS-1 cells. J Biol Chem 279, 44370–44375. 10.1074/jbc.M311842200.

21. Cline, G.W., Pongratz, R.L., Zhao, X., and Papas, K.K. (2011). Rates of insulin secretion in INS-1 cells are enhanced by coupling to anaplerosis and Kreb’s cycle flux independent of ATP synthesis. Biochem Biophys Res Commun 415, 30–35. 10.1016/j.bbrc.2011.09.153.

22. Simpson, N.E., Khokhlova, N., Oca-Cossio, J.A., and Constantinidis, I. (2006). Insights into the role of anaplerosis in insulin secretion: A 13C NMR study. Diabetologia 49, 1338–1348. 10.1007/s00125-006-0216-5.

23. Malinowski, R.M., Ghiasi, S.M., Mandrup-Poulsen, T., Meier, S., Lerche, M.H., Ardenkjær-Larsen, J.H., and Jensen, P.R. (2020). Pancreatic β-cells respond to fuel pressure with an early metabolic switch. Scientific Reports 10. 10.1038/s41598-020-72348-1.

24. Benninger, R.K.P., and Hodson, D.J. (2018). New Understanding of β-Cell Heterogeneity and In Situ Islet Function. Diabetes 67, 537-547.

25. Benninger, R.K.P., and Kravets, V. (2021). The physiological role of β-cell heterogeneity in pancreatic islet function. Nature Reviews Endocrinology. 10.1038/s41574-021-00568-0.

26. Nasteska, D., Fine, N.H.F., Ashford, F.B., Cuozzo, F., Viloria, K., Smith, G., Dahir, A., Dawson, P.W.J., Lai, Y.-C., Bastidas-Ponce, A., et al. (2021). PDX1LOW MAFALOW β-cells contribute to islet function and insulin release. Nature Communications 12, 674. 10.1038/s41467-020-20632-z.

27. Rodriguez-Diaz, R., Dando, R., Jacques-Silva, M.C., Fachado, A., Molina, J., Abdulreda, M.H., Ricordi, C., Roper, S.D., Berggren, P.O., and Caicedo, A. (2011). Alpha cells secrete acetylcholine as a non-neuronal paracrine signal priming beta cell function in humans. Nature Medicine 17, 888–892. 10.1038/nm.2371.

28. Cabrera, O., Berman, D.M., Kenyon, N.S., Ricordi, C., Berggren, P.O., and Caicedo, A. (2006). The unique cytoarchitecture of human pancreatic islets has implications for islet cell function. Proceedings of the National Academy of Sciences of the United States of America 103, 2334–2339. 0510790103 [pii]10.1073/pnas.0510790103.

29. Hodson, D.J., Mitchell, R.K., Bellomo, E.A., Sun, G., Vinet, L., Meda, P., Li, D., Li, W.H., Bugliani, M., Marchetti, P., et al. (2013). Lipotoxicity disrupts incretin-regulated human beta cell connectivity. Journal of Clinical Investigation 123, 4182–4194. 10.1172/JCI68459.

30. Spegel, P., Sharoyko, V.V., Goehring, I., Danielsson, A.P., Malmgren, S., Nagorny, C.L., Andersson, L.E., Koeck, T., Sharp, G.W., Straub, S.G., et al. (2013). Time-resolved metabolomics analysis of beta-cells implicates the pentose phosphate pathway in the control of insulin release. Biochem J 450, 595–605. 10.1042/BJ20121349.

31. Wallace, M., Whelan, H., and Brennan, L. (2013). Metabolomic analysis of pancreatic beta cells following exposure to high glucose. Biochimica et Biophysica Acta (BBA) - General Subjects 1830, 2583–2590. 10.1016/j.bbagen.2012.10.025.

32. Barsby, T., Vähäkangas, E., Ustinov, J., Montaser, H., Ibrahim, H., Lithovius, V., Kuuluvainen, E., Chandra, V., Saarimäki-Vire, J., Katajisto, P., et al. (2023). Aberrant metabolite trafficking and fuel sensitivity in human pluripotent stem cell-derived islets. Cell Reports 42. 10.1016/j.celrep.2023.112970.

33. Balboa, D., Barsby, T., Lithovius, V., Saarimaki-Vire, J., Omar-Hmeadi, M., Dyachok, O., Montaser, H., Lund, P.E., Yang, M., Ibrahim, H., et al. (2022). Functional, metabolic and transcriptional maturation of human pancreatic islets derived from stem cells. Nat Biotechnol 40, 1042–1055. 10.1038/s41587-022-01219-z.

34. Lu, Mulder, H., Zhao, P., Burgess, S.C., Jensen, M.V., Kamzolova, S., Newgard, C.B., and Sherry, A.D. (2002). 13C NMR isotopomer analysis reveals a connection between pyruvate cycling and glucose-stimulated insulin secretion (GSIS). Proc Natl Acad Sci U S A 99, 2708–2713. 10.1073/pnas.052005699.

35. Alonso, L., Piron, A., Morán, I., Guindo-Martínez, M., Bonàs-Guarch, S., Atla, G., Miguel-Escalada, I., Royo, R., Puiggròs, M., Garcia-Hurtado, X., et al. (2021). TIGER: The gene expression regulatory variation landscape of human pancreatic islets. Cell Reports 37. 10.1016/j.celrep.2021.109807.

36. Goldberg, E., Eddy, E.M., Duan, C., and Odet, F. (2009). LDHC: The Ultimate Testis-Specific Gene. Journal of Andrology 31, 86–94. 10.2164/jandrol.109.008367.

37. Segerstolpe, Å., Palasantza, A., Eliasson, P., Andersson, E.-M., Andréasson, A.-C., Sun, X., Picelli, S., Sabirsh, A., Clausen, M., Bjursell, M.K., et al. (2016). Single-Cell Transcriptome Profiling of Human Pancreatic Islets in Health and Type 2 Diabetes. Cell Metabolism 24, 593–607. 10.1016/j.cmet.2016.08.020.

38. Uhlen, M., Fagerberg, L., Hallstrom, B.M., Lindskog, C., Oksvold, P., Mardinoglu, A., Sivertsson, A., Kampf, C., Sjostedt, E., Asplund, A., et al. (2015). Proteomics. Tissue-based map of the human proteome. Science 347, 1260419. 10.1126/science.1260419.

39. Thul, P.J., Akesson, L., Wiking, M., Mahdessian, D., Geladaki, A., Ait Blal, H., Alm, T., Asplund, A., Bjork, L., Breckels, L.M., et al. (2017). A subcellular map of the human proteome. Science 356. 10.1126/science.aal3321.

40. van Gurp, L., Fodoulian, L., Oropeza, D., Furuyama, K., Bru-Tari, E., Vu, A.N., Kaddis, J.S., Rodriguez, I., Thorel, F., and Herrera, P.L. (2022). Generation of human islet cell type-specific identity genesets. Nat Commun 13, 2020. 10.1038/s41467-022-29588-8.

41. Pullen, T.J., Sylow, L., Sun, G., Halestrap, A.P., Richter, E.A., and Rutter, G.A. (2012). Overexpression of Monocarboxylate Transporter-1 (Slc16a1) in Mouse Pancreatic beta-Cells Leads to Relative Hyperinsulinism During Exercise. Diabetes 61, 1719–1725. 10.2337/db11-1531.

42. Zhao, C., Wilson, M.C., Schuit, F., Halestrap, A.P., and Rutter, G.A. (2001). Expression and distribution of lactate/monocarboxylate transporter isoforms in pancreatic islets and the exocrine pancreas. Diabetes 50, 361–366.

43. Deng, H., Gao, Y., Trappetti, V., Hertig, D., Karatkevich, D., Losmanova, T., Urzi, C., Ge, H., Geest, G.A., Bruggmann, R., et al. (2022). Targeting lactate dehydrogenase B-dependent mitochondrial metabolism affects tumor initiating cells and inhibits tumorigenesis of non-small cell lung cancer by inducing mtDNA damage. Cellular and Molecular Life Sciences 79. 10.1007/s00018-022-04453-5.

44. Ždralević, M., Brand, A., Di Ianni, L., Dettmer, K., Reinders, J., Singer, K., Peter, K., Schnell, A., Bruss, C., Decking, S.-M., et al. (2018). Double genetic disruption of lactate dehydrogenases A and B is required to ablate the “Warburg effect” restricting tumor growth to oxidative metabolism. Journal of Biological Chemistry 293, 15947–15961. 10.1074/jbc.RA118.004180.

45. Jekabsons, M., San Martín, A., Ceballo, S., Ruminot, I., Lerchundi, R., Frommer, W.B., and Barros, L.F. (2013). A Genetically Encoded FRET Lactate Sensor and Its Use To Detect the Warburg Effect in Single Cancer Cells. PLoS ONE 8. 10.1371/journal.pone.0057712.

46. Sdao, S.M., Ho, T., Poudel, C., Foster, H.R., De Leon, E.R., Adams, M.T., Lee, J.H., Blum, B., Rane, S.G., and Merrins, M.J. (2021). CDK2 limits the highly energetic secretory program of mature beta cells by restricting PEP cycle-dependent K(ATP) channel closure. Cell Rep 34, 108690. 10.1016/j.celrep.2021.108690.

47. Shibata, S., Sogabe, S., Miwa, M., Fujimoto, T., Takakura, N., Naotsuka, A., Kitamura, S., Kawamoto, T., and Soga, T. (2021). Identification of the first highly selective inhibitor of human lactate dehydrogenase B. Scientific Reports 11. 10.1038/s41598-021-00820-7.

48. Blanchi, B., Taurand, M., Colace, C., Thomaidou, S., Audeoud, C., Fantuzzi, F., Sawatani, T., Gheibi, S., Sabadell-Basallote, J., Boot, F.W.J., et al. (2023). EndoC-βH5 cells are storable and ready-to-use human pancreatic beta cells with physiological insulin secretion. Molecular Metabolism 76. 10.1016/j.molmet.2023.101772.

49. Schuit, F., De Vos, A., Farfari, S., Moens, K., Pipeleers, D., Brun, T., and Prentki, M. (1997). Metabolic fate of glucose in purified islet cells. Glucose-regulated anaplerosis in beta cells. J Biol Chem 272, 18572–18579. 10.1074/jbc.272.30.18572.

50. Lemaire, K., Thorrez, L., and Schuit, F. (2016). Disallowed and Allowed Gene Expression: Two Faces of Mature Islet Beta Cells. Annu Rev Nutr 36, 45–71. 10.1146/annurev-nutr-071715-050808.

51. Sato, Y., Endo, H., Okuyama, H., Takeda, T., Iwahashi, H., Imagawa, A., Yamagata, K., Shimomura, I., and Inoue, M. (2011). Cellular hypoxia of pancreatic beta-cells due to high levels of oxygen consumption for insulin secretion in vitro. J Biol Chem 286, 12524–12532. 10.1074/jbc.M110.194738.

52. Bensellam, M., Duvillie, B., Rybachuk, G., Laybutt, D.R., Magnan, C., Guiot, Y., Pouyssegur, J., and Jonas, J.C. (2012). Glucose-induced O(2) consumption activates hypoxia inducible factors 1 and 2 in rat insulin-secreting pancreatic beta-cells. PLoS One 7, e29807. 10.1371/journal.pone.0029807.

53. Taddeo, E.P., Stiles, L., Sereda, S., Ritou, E., Wolf, D.M., Abdullah, M., Swanson, Z., Wilhelm, J., Bellin, M., McDonald, P., et al. (2018). Individual islet respirometry reveals functional diversity within the islet population of mice and human donors. Mol Metab 16, 150–159. 10.1016/j.molmet.2018.07.003.

54. Sanchez, P.K.M., Khazaei, M., Gatineau, E., Geravandi, S., Lupse, B., Liu, H., Dringen, R., Wojtusciszyn, A., Gilon, P., Maedler, K., and Ardestani, A. (2021). LDHA is enriched in human islet alpha cells and upregulated in type 2 diabetes. Biochem Biophys Res Commun 568, 158–166. 10.1016/j.bbrc.2021.06.065.

55. Moin, A.S.M., Cory, M., Gurlo, T., Saisho, Y., Rizza, R.A., Butler, P.C., and Butler, A.E. (2020). Pancreatic alpha-cell mass across adult human lifespan. Eur J Endocrinol 182, 219–231. 10.1530/EJE-19-0844.

56. Zaborska, K.E., Dadi, P.K., Dickerson, M.T., Nakhe, A.Y., Thorson, A.S., Schaub, C.M., Graff, S.M., Stanley, J.E., Kondapavuluru, R.S., Denton, J.S., and Jacobson, D.A. (2020). Lactate activation of alpha-cell KATP channels inhibits glucagon secretion by hyperpolarizing the membrane potential and reducing Ca(2+) entry. Mol Metab 42, 101056. 10.1016/j.molmet.2020.101056.

57. Pullen, T.J., and Rutter, G.A. (2013). When less is more: the forbidden fruits of gene repression in the adult beta-cell. Diabetes Obes Metab 15, 503–512. 10.1111/dom.12029.

58. Fu, A., Alvarez-Perez, J.C., Avizonis, D., Kin, T., Ficarro, S.B., Choi, D.W., Karakose, E., Badur, M.G., Evans, L., Rosselot, C., et al. (2020). Glucose-dependent partitioning of arginine to the urea cycle protects β-cells from inflammation. Nature Metabolism 2, 432–446. 10.1038/s42255-020-0199-4.

59. Ainscow, E.K., and Rutter, G.A. (2002). Glucose-Stimulated Oscillations in Free Cytosolic ATP Concentration Imaged in Single Islet β-Cells. Diabetes 51, S162–S170. 10.2337/diabetes.51.2007.S162.

60. Campbell, J.E., and Newgard, C.B. (2021). Mechanisms controlling pancreatic islet cell function in insulin secretion. Nat Rev Mol Cell Biol 22, 142–158. 10.1038/s41580-020-00317-7.

61. Gerber, P.A., and Rutter, G.A. (2017). The Role of Oxidative Stress and Hypoxia in Pancreatic Beta-Cell Dysfunction in Diabetes Mellitus. Antioxid Redox Signal 26, 501–518. 10.1089/ars.2016.6755.

62. Bensellam, M., Jonas, J.C., and Laybutt, D.R. (2018). Mechanisms of beta-cell dedifferentiation in diabetes: recent findings and future research directions. J Endocrinol 236, R109–R143. 10.1530/JOE-17-0516.

63. Marselli, L., Piron, A., Suleiman, M., Colli, M.L., Yi, X., Khamis, A., Carrat, G.R., Rutter, G.A., Bugliani, M., Giusti, L., et al. (2020). Persistent or Transient Human β Cell Dysfunction Induced by Metabolic Stress: Specific Signatures and Shared Gene Expression with Type 2 Diabetes. Cell Reports 33. 10.1016/j.celrep.2020.108466.

64. Viloria, K., Nasteska, D., Ast, J., Hasib, A., Cuozzo, F., Heising, S., Briant, L.J.B., Hewison, M., and Hodson, D.J. (2023). GC-Globulin/Vitamin D–Binding Protein Is Required for Pancreatic α-Cell Adaptation to Metabolic Stress. Diabetes 72, 275–289. 10.2337/db22-0326.

65. Fernandez, C.A., Des Rosiers, C., Previs, S.F., David, F., and Brunengraber, H. (1996). Correction of 13C mass isotopomer distributions for natural stable isotope abundance. J Mass Spectrom 31, 255–262. 10.1002/(SICI)1096-9888(199603)31:3<255::AID-JMS290>3.0.CO;2-3.

66. Kazimierczuk, K., and Orekhov, V.Y. (2011). Accelerated NMR Spectroscopy by Using Compressed Sensing. Angewandte Chemie International Edition 50, 5556–5559. 10.1002/anie.201100370.

67. Delaglio, F., Grzesiek, S., Vuister, G., Zhu, G., Pfeifer, J., and Bax, A. (1995). NMRPipe: A multidimensional spectral processing system based on UNIX pipes. Journal of Biomolecular NMR 6. 10.1007/bf00197809.

68. Ludwig, C., and Günther, U.L. (2011). MetaboLab - advanced NMR data processing and analysis for metabolomics. BMC Bioinformatics 12. 10.1186/1471-2105-12-366.

69. Tantama, M., Martinez-Francois, J.R., Mongeon, R., and Yellen, G. (2013). Imaging energy status in live cells with a fluorescent biosensor of the intracellular ATP-to-ADP ratio. Nat Commun 4, 2550. 10.1038/ncomms3550.

70. Gavet, O., and Pines, J. (2010). Progressive activation of CyclinB1-Cdk1 coordinates entry to mitosis. Dev Cell 18, 533–543. 10.1016/j.devcel.2010.02.013.

71. Viloria, K., Nasteska, D., Briant, L.J.B., Heising, S., Larner, D.P., Fine, N.H.F., Ashford, F.B., da Silva Xavier, G., Ramos, M.J., Hasib, A., et al. (2020). Vitamin-D-Binding Protein Contributes to the Maintenance of α Cell Function and Glucagon Secretion. Cell Reports 31. 10.1016/j.celrep.2020.107761.

72. Satija, R., Farrell, J.A., Gennert, D., Schier, A.F., and Regev, A. (2015). Spatial reconstruction of single-cell gene expression data. Nat Biotechnol 33, 495–502. 10.1038/nbt.3192.

73. Aibar, S., Gonzalez-Blas, C.B., Moerman, T., Huynh-Thu, V.A., Imrichova, H., Hulselmans, G., Rambow, F., Marine, J.C., Geurts, P., Aerts, J., et al. (2017). SCENIC: single-cell regulatory network inference and clustering. Nat Methods 14, 1083–1086. 10.1038/nmeth.4463.

74. Arda, H.E., Li, L., Tsai, J., Torre, E.A., Rosli, Y., Peiris, H., Spitale, R.C., Dai, C., Gu, X., Qu, K., et al. (2016). Age-Dependent Pancreatic Gene Regulation Reveals Mechanisms Governing Human beta Cell Function. Cell Metab 23, 909–920. 10.1016/j.cmet.2016.04.002.

75. Bray, N.L., Pimentel, H., Melsted, P., and Pachter, L. (2016). Near-optimal probabilistic RNA-seq quantification. Nat Biotechnol 34, 525–527. 10.1038/nbt.3519.

76. Akerman, I., Kasaai, B., Bazarova, A., Sang, P.B., Peiffer, I., Artufel, M., Derelle, R., Smith, G., Rodriguez-Martinez, M., Romano, M., et al. (2020). A predictable conserved DNA base composition signature defines human core DNA replication origins. Nat Commun 11, 4826. 10.1038/s41467-020-18527-0.

77. Baron, M., Veres, A., Wolock, S.L., Faust, A.L., Gaujoux, R., Vetere, A., Ryu, J.H., Wagner, B.K., Shen-Orr, S.S., Klein, A.M., et al. (2016). A Single-Cell Transcriptomic Map of the Human and Mouse Pancreas Reveals Inter-and Intra-cell Population Structure. Cell Syst 3, 346–360 e344. 10.1016/j.cels.2016.08.011.

78. Xin, Y., Kim, J., Okamoto, H., Ni, M., Wei, Y., Adler, C., Murphy, A.J., Yancopoulos, G.D., Lin, C., and Gromada, J. (2016). RNA Sequencing of Single Human Islet Cells Reveals Type 2 Diabetes Genes. Cell Metab 24, 608–615. 10.1016/j.cmet.2016.08.018.

79. Mawla, A.M., and Huising, M.O. (2019). Navigating the Depths and Avoiding the Shallows of Pancreatic Islet Cell Transcriptomes. Diabetes 68, 1380–1393. 10.2337/dbi18-0019.

80. Lawlor, N., George, J., Bolisetty, M., Kursawe, R., Sun, L., Sivakamasundari, V., Kycia, I., Robson, P., and Stitzel, M.L. (2017). Single-cell transcriptomes identify human islet cell signatures and reveal cell-type–specific expression changes in type 2 diabetes. Genome Research 27, 208–222. 10.1101/gr.212720.116.

81. Wang, Y.J., Schug, J., Won, K.J., Liu, C., Naji, A., Avrahami, D., Golson, M.L., and Kaestner, K.H. (2016). Single-Cell Transcriptomics of the Human Endocrine Pancreas. Diabetes 65, 3028–3038. 10.2337/db16-0405.

82. Xin, Y., Gutierrez, G.D., Okamoto, H., Kim, J., Lee, A.H., Adler, C., Ni, M., Yancopoulos, G.D., Murphy, A.J., and Gromada, J. (2018). Pseudotime Ordering of Single Human beta-Cells Reveals States of Insulin Production and Unfolded Protein Response. Diabetes. 10.2337/db18-0365.

83. Enge, M., Arda, H.E., Mignardi, M., Beausang, J., Bottino, R., Kim, S.K., and Quake, S.R. (2017). Single-Cell Analysis of Human Pancreas Reveals Transcriptional Signatures of Aging and Somatic Mutation Patterns. Cell 171, 321–330 e314. 10.1016/j.cell.2017.09.004.

84. Akerman, I., Tu, Z., Beucher, A., Rolando, D.M.Y., Sauty-Colace, C., Benazra, M., Nakic, N., Yang, J., Wang, H., Pasquali, L., et al. (2017). Human Pancreatic beta Cell lncRNAs Control Cell-Specific Regulatory Networks. Cell Metab 25, 400–411. 10.1016/j.cmet.2016.11.016.

