## Supplementary text and figures for "High-resolution glucose tracing and in situ imaging reveals regulated lactate production in human pancreatic β cells"

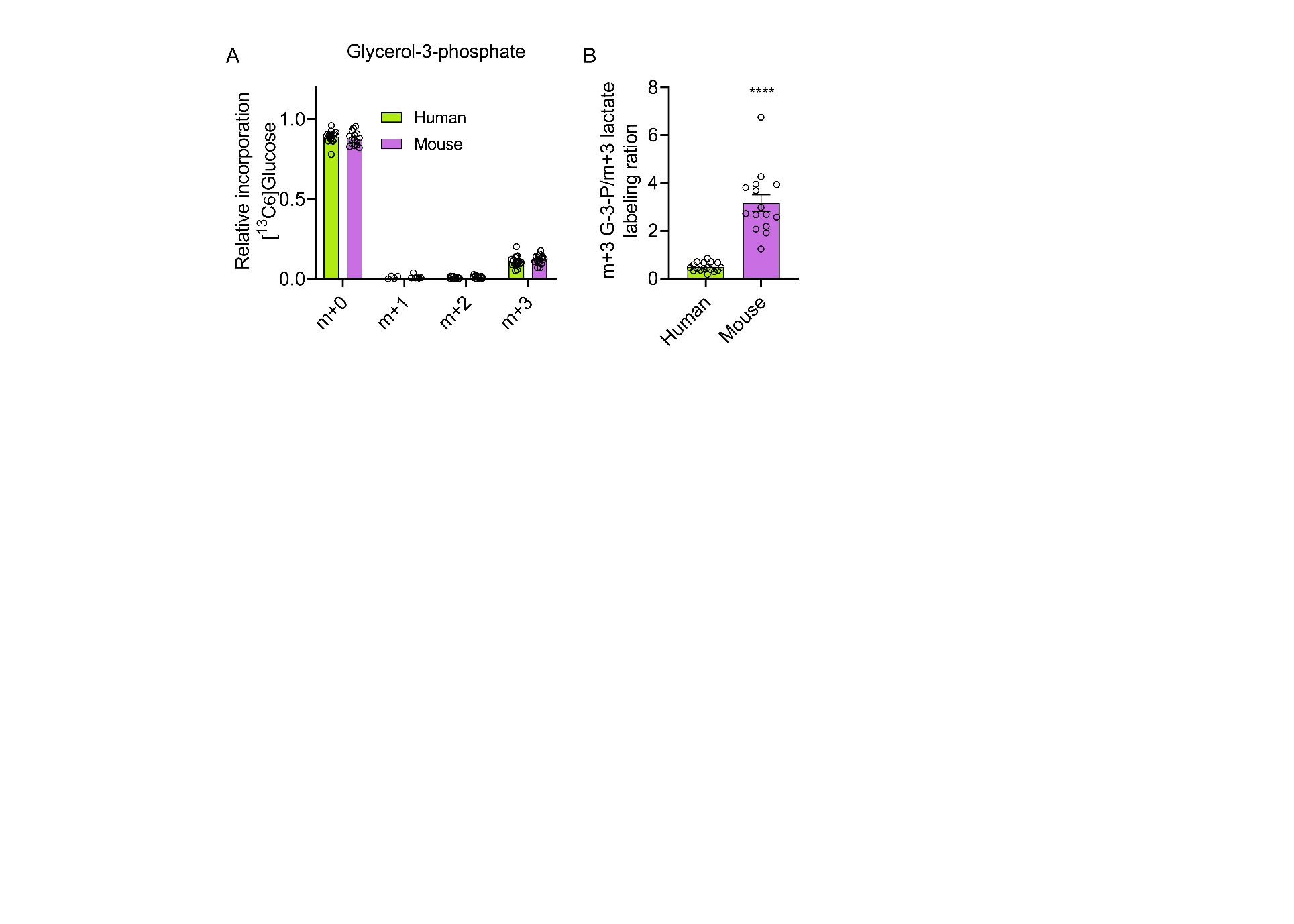

**Figure S1: Glycolytic contribution to the accumulation of lactate. A)** MID analysis showing similar incorporation of ^13^C from ^13^C_6_ glucose into m+0 and m+3 glycerol-3-phosphate in human *versus* mouse islets (two-way ANOVA and Sidak’s post-hoc test ) **B)** Labeling ratio of fully labelled G-3-P over fully-labeled lactate is significantly higher in mouse than human islets (unpaired t-test). For all data, n = 17 islet preparations, 9 donors and n = 15 islet preparations, 8 animals. Bar graphs show individual datapoints and mean ± SEM.

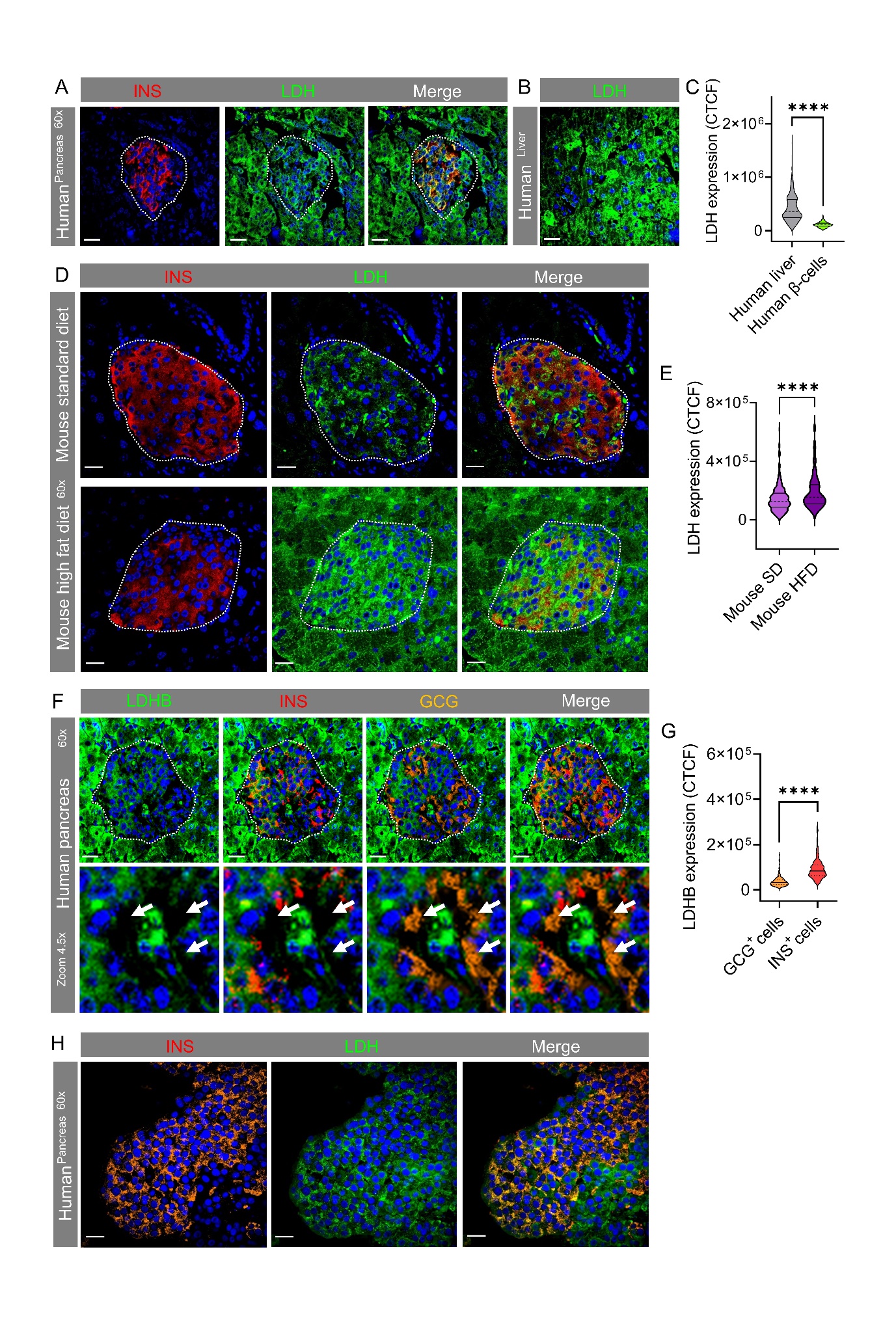

**Figure S2: Higher magnification re-quantification of LDH and LDHB in human and mouse pancreas sections. A-C)** New sections were cut from the same donor pancreata (A) and livers (B) as used in Figure 2E and I, before repeat immunostaining for LDH and quantification with a 60x, 1.30 NA objective. Note the much higher LDH expression in human liver versus β-cells (C), identified by insulin positivity (n = 300 cells, 3 donors). **D, E)** Pancreata from mice fed either standard diet or high fat diet were re-imaged with a 60x, 1.30 NA objective, showing similar results to those in Figure 2F (D) (n = 279-295 cells, 3 animals). **F, G**) New sections were cut from the same donor pancreata (A) as used in Figure 4A, before repeat immunostaining for LDHB and quantification with a 60x, 1.30 NA objective. As previously shown, LHDB is expressed at very low to undetectable levels in α-cells (n= 300 cells, 3 donors). **H)** LDH (LDHA + LDHB) immunostaining in isolated human islets, showing a similar pattern and intensity to that seen in pancreatic slices (Figure 6). All data were analyzed using Mann-Whitney test. Scale bar = 30 µm. Violin plot shows median and interquartile range.

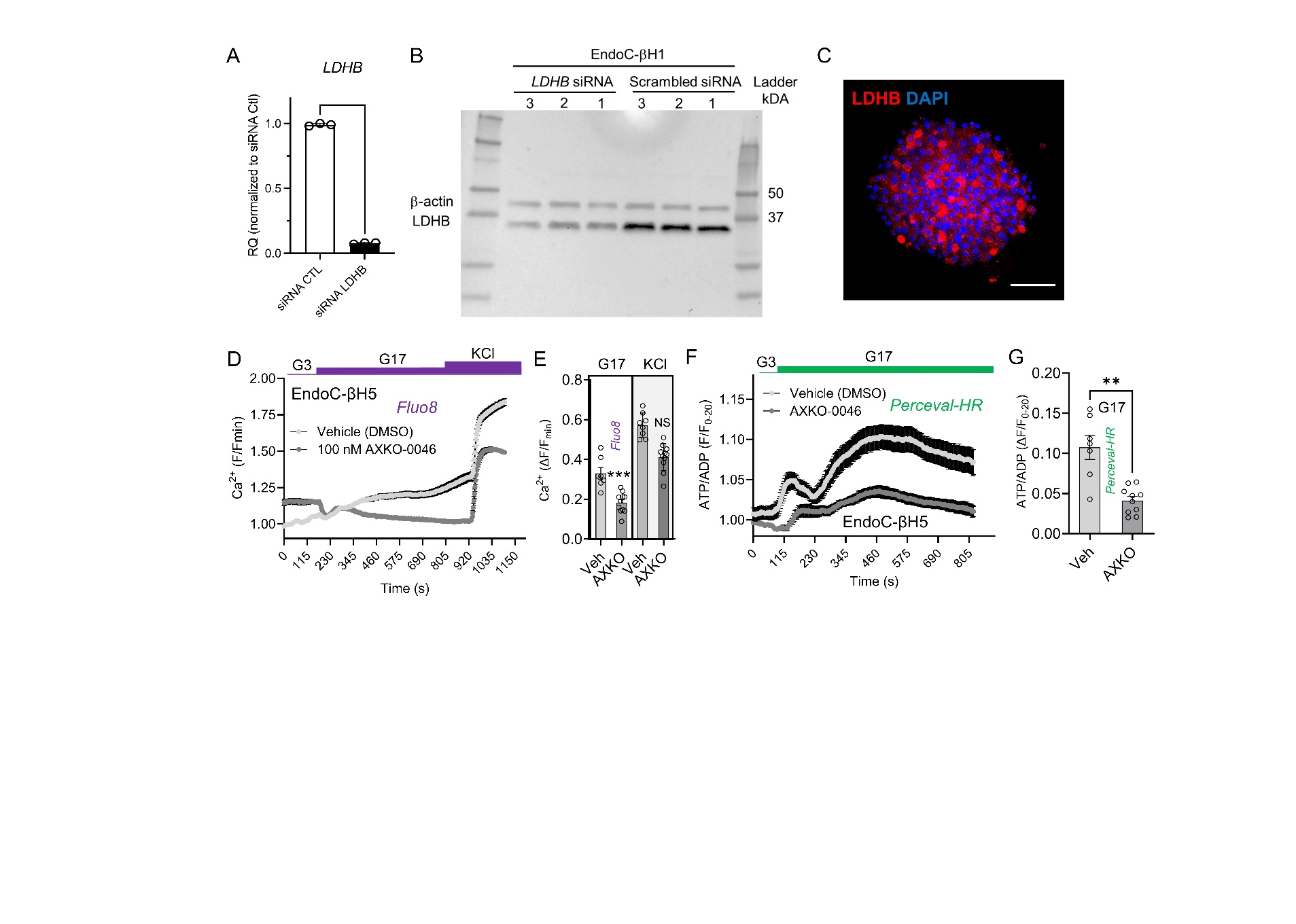

**Figure S3:** LDHB knockdown, LDHB expression and LDHB inhibition in EndoC-βH1 and EndoC-βH5 cells. **A)** qPCR analysis of *LDHB* expression in RNA extracted from EndoC-βH1 cells transfected with siRNA CTL (scrambled) and siRNA *LDHB*. Values were normalized to a house keeping gene, *PPIA*, followed by normalization to siRNA CTL. **B)** Protein lysates were extracted from EndoC-βH1 cells transfected with LDHB and CTL siRNA. Expected band for LDHB is ~35kDa. β-actin is used as loading control with an expected band of ~45kDa. (n=3 is used for both experimental groups; full uncropped blot is shown). **C)** EndoC-βH5 spheroid immunostained for LDHB (n = 5 spheroids) (scale bar 53 µm). **D, E)** Vehicle-treated EndoC-βH5 spheroids respond to 17 mM glucose with increases in cytosolic Ca^2+^, an effect suppressed by pre-incubation with 100 nM AXKO-0046, as shown by mean ± SEM traces (D) and summary bar graph (E) (n = 11-13 spheroids) (Mann-Whitney test). **F, G)** As for D, E), but ATP/ADP ratios measure using Perceval-HR (n = 7-10 spheroids) (Mann-Whitney test). Bar graphs show individual datapoints and mean ± SEM. G3, 3 mM glucose; G17, 17 mM glucose.

**Table S1: Human islet donor characteristics.** BMI, body mass index. IFG, impaired fasting glucose.

| **Unique identifier** | **Age group (years)** | **Gender** | **BMI (Kg/m^2^)** | **Glycemia (mmol/L)* HbA1C (%)** | **History of diabetes**** | **Islet purity (%)** | **Islet culture duration (h)** | **Country of origin** |
| --- | --- | --- | --- | --- | --- | --- | --- | --- |
| **HP1404** | 50-55 | ♂ | 29.4 | 7.8 mmol/L | N/A | 80 | 18 | Italy |
| **HP1406** | 60-65 | ♂ | 26.1 | N/A | N/A | 90 | 96 | Italy |
| **HP1408** | 55-60 | ♀ | 19.0 | N/A | N/A | 90 | 18 | Italy |
| **HP1416** | 60-65 | ♂ | 31.1 | N/A | No but IFG | 75 | 20 | Italy |
| **HP1419** | 55-60 | ♂ | 22.8 | 7.3 mmol/L | N/A | 90 | 18 | Italy |
| **HP1431** | 60-65 | ♀ | 26.9 | 8.0 mmol/L | N/A | 90 | 18 | Italy |
| **HI1117** | 45-50 | ♂ | 24.0 | 5.4% | N/A | 80 | N/A | France |
| **HI1120** | 50-55 | ♂ | 29.5 | 5.7% | N/A | 90 | N/A | France |
| **HI1121** | 60-65 | ♂ | 32 | 5.5% | N/A | 90 | N/A | France |
| **R496** | 68 | ♀ | 22.3 | 5.6% | N/A | 95 | N/A | Canada |
| **HP2337** | 36 | ♀ | 29.71 |  | N/A | 80 | N/A | UK |
| **HP2338** | 51 | ♀ | 30.11 |  | N/A | 70 | N/A | UK |
| **H1236** | 49 | ♂ | 29.2 | 5.7% | N/A | 90 | N/A | France |
| **HP2339** | 39 | ♂ | 30.39 |  | N/A | 70 | N/A | UK |
| **R511** | 47 | ♀ | 19.2 | 5.9% | N/A | 90 | N/A | Canada |
| **R512** | 63 | ♂ | 34.7 | 5.9% | N/A | 90 | N/A | Canada |
| **R513** | 62 | ♀ | 26.4 | 5.3% | N/A | 40 | N/A | Canada |

**Table S2: Retention times, ion counts and MID data for mouse islet GC-MS.**

**Table S3: Retention times, ion counts and MID data for human islet GC-MS.**

**Table S4: Human pancreas and liver donor characteristics.** BMI, body mass index. ND, non-diabetic. NASH, non-alcoholic steatohepatitis.

| **Sample ID** | **Disease** | **Tissue** | **Age** | **BMI (Kg/m^2^)** | **Sex** | **Cause of death** |
| --- | --- | --- | --- | --- | --- | --- |
| 43-D | ND | Adult pancreas | 78 | - | F | Myocardial infarction |
| 137-D | ND | Adult pancreas | 67 | - | M | Ischaemic heart disease |
| 77-C | ND | Adult pancreas | 56 | - | F | Cerebral aneurysm |
| LT6 T0 | NASH | Adult liver | 54 | 28.4 |  | Stroke |
| LT7 T12 | NASH | Adult liver | 41 | 35 |  | Intracranial haemorrhage |
| LT8 T48 | NASH | Adult liver | 69 | 24.8 |  | Intracranial haemorrhage |

**Table S5: Markers used for cell type selection in UMAP plots**

| **Cell type** | **Markers** |
| --- | --- |
| Endothelial | CD34, PECAM1, CD93 |
| Proliferative | MKI67 |
| Mesenchymal | TNFAIP6, THY1 |
| Acinar | CTRC, PRSS1 |
| Ductal | CFTR, KRT19, SOX9 |
| Antigen Presenting | HLA-DRA, CXCL8, HLA-DRB1, CD83 |
| Beta | PDX1, INS, IAPP |
| Delta | SST, PDX1 |
| Alpha | GCG |
| Epsilon | GHRL |
| PP | PYY |
| Non-endocrine cells | CHGA-, NEUROD1- |
| Endocrine | CHGA+, NEUROD1+ |
